# A GATA2-PROX1-FOXC2 regulatory axis integrates shear stress signaling with lymphatic vascular development

**DOI:** 10.64898/2026.06.25.734612

**Authors:** Xin Geng, Md. Riaj Mahamud, Wojciech Rosikiewicz, Martijn A. van der Ent, Scott D Zawieja, Hong Chen, Audrey C.A. Cleuren, Chunliang Li, Michael J Davis, R. Sathish Srinivasan

## Abstract

The two-hit hypothesis provides a novel framework for incompletely penetrant vascular disorders. Emberger syndrome, caused by heterozygous loss-of-function mutations in mechanosensitive transcription factor (TF) GATA2, is associated with lymphedema in a subset of patients, suggesting that modifiers influence disease development. Heterozygous loss-of-function mutations in mechanosensitive TF FOXC2 also cause lymphedema. Complete deletion of either factor in lymphatic endothelial cells (LECs) causes severe, overlapping defects, including complete loss of lymphatic valves, whereas single heterozygous mutants exhibit milder phenotypes, suggesting genetic buffering. However, whether GATA2 and FOXC2 interact with each other and with shear stress-dependent transcriptional programs remains unclear. Here, we show that *Gata2^+/-^;Foxc2^+/-^* mice develop profound lymphatic vascular defects, including absent lymphatic valves, perinatal lethality, and valve dysfunction, revealing dosage-dependent cooperation between GATA2 and FOXC2. Using a Cre-dependent model of LEC-specific GATA2 overexpression, we found that increased GATA2 dosage downregulated PROX1 and disrupted lymphatic vascular development. ATAC-seq and bulk RNA-seq of shear-exposed human LECs showed that PROX1 governs shear-responsive genes, including *KLF2* and *KLF4,* whereas FOXC2 modulates PROX1-dependent and independent gene subsets. Together, these findings identify PROX1 as a central integrator of shear stress-responsive transcriptional programs and reveal that balanced GATA2 and FOXC2 dosage preserves this network during lymphatic vascular development.

## INTRODUCTION

The mammalian lymphatic vasculature is composed of lymphatic capillaries, collecting lymphatic vessels, lymphatic valves (LVs), and lymphovenous valves (LVVs) ^1–3^. Together, these structures collect and transport interstitial fluid, dietary lipids, and immune cells from tissues to lymph nodes and ultimately back to the bloodstream. Proper lymphatic vascular function is essential for preventing tissue swelling, enabling lipid absorption, and supporting immune responses. Accordingly, impaired lymphatic vascular development or function causes lymphedema, immune dysfunction, fibrosis, and other debilitating disorders.

Lymphatic vascular damage can arise from developmental abnormalities, surgical procedures, or infections ^3,4^. Currently, there are no effective therapies to repair damaged lymphatic vessels or valves. Thus, a deeper understanding of the mechanisms governing lymphatic vascular development and disease is essential for developing new therapeutic strategies for currently incurable lymphatic disorders.

A major unresolved question is why heterozygous mutations in lymphatic disease genes often produce variable phenotypes, ranging from clinically silent defects to severe congenital disease. This incomplete penetrance suggests that additional genetic or environmental modifiers may determine whether partial loss of a single regulatory factor results in disease onset. The two-hit hypothesis, originally developed to explain tumor onset in patients carrying heterozygous mutations in tumor suppressor genes, is increasingly being considered as a framework for incompletely penetrant vascular disorders ^5–10^.

The zinc finger transcription factor (TF) GATA2 is essential for the development of multiple cell types, including hematopoietic and endothelial cells. Heterozygous mutations in both the coding and non-coding regions of GATA2 have been identified in patients with Emberger syndrome, which is characterized by myelodysplastic syndrome (MDS) and acute myeloid leukemia (AML) ^11,12^. Approximately 25% of these patients also develop lymphedema, often preceding the onset of hematologic disease ^13^. The timing of lymphedema onset is highly variable, ranging from in utero to adulthood. Similarly, the severity of the lymphatic disorder can vary widely, from mild edema to bilateral limb lymphedema and even hydrops fetalis. The underlying reasons for this variable penetrance remain unknown, but these observations suggest that GATA2 haploinsufficiency may require additional modifiers to produce overt lymphatic disease.

We and others have shown that GATA2 is expressed in all lymphatic endothelial cells (LECs), with the highest levels in LVs and LVVs ^12,14–16^. Conditional deletion of both *Gata2* alleles in the lymphatic vasculature of mice leads to the complete absence of LVs and LVVs, along with dilated lymphatic vessels. Although valvular endothelial cells are specified in GATA2-deficient mice, they are unable to survive and undergo morphogenesis into valves ^16^. *Gata2^+/-^* mice have been shown to exhibit dilated thoracic duct and impaired regrowth of lymphatic vessels following lymphadenectomy ^15,17^. However, these findings do not fully account for the development of congenital lymphedema in patients. Whether LVs or LVVs are defective in patients or *Gata2^+/-^* mice remains unknown.

Several putative downstream targets of GATA2 have been proposed, including PROX1, FOXC2, miR-126, ANGPT2, FLT4, and FAT4 ^12,14–16,18^. GATA2 also interacts with and stabilizes the transcriptional regulator MDFIC ^19^. Many of these GATA2 targets are themselves implicated in primary lymphedema, raising the possibility that GATA2 functions within a dosage-sensitive regulatory network. Among them, the relationship of GATA2 with the TF FOXC2 is particularly compelling.

Mutations in FOXC2 cause lymphedema-distichiasis syndrome, which is characterized by late-onset lymphedema with high penetrance ^20^. FOXC2 is expressed in all LECs, with enrichment in valvular endothelial cells, mirroring the expression of GATA2^21^. Functionally, FOXC2 is essential for LV and LVV morphogenesis. Mice lacking *Foxc2* fail to form LVs and LVVs, and *Foxc2^+/-^* mice show a reduced number of valves ^21–24^. Thus, GATA2 and FOXC2 are expressed in overlapping lymphatic vascular compartments, are both required for valve development, and are each linked to human lymphedema. These observations raise the possibility that partial loss of both factors may compromise a shared regulatory program required for lymphatic vascular development.

Despite substantial overlap in expression and activity, the functional relationship between GATA2 and FOXC2 remains incompletely defined. Chromatin immunoprecipitation assays have shown that GATA2 and FOXC2 can co-occupy regulatory elements of numerous genes, including a critical *PROX1* enhancer ^25^. However, how FOXC2 and GATA2 regulate PROX1 expression remains unclear. In particular, whether these factors act redundantly, hierarchically, or in a dosage-sensitive manner to control PROX1 expression and lymphatic vascular development is not known. Addressing this question is important for understanding why partial loss of either GATA2 or FOXC2 produces relatively mild phenotypes in mice, yet predisposes to lymphedema in humans.

To address this knowledge gap in vivo, we systematically analyzed LVs, LVVs, and lymphatic vessel development of in *Gata2^+/-^*, *Foxc2^+/-^* and *Gata2^+/-^*;*Foxc2^+/-^* mice, and quantified LV function. We also employed a novel Cre-dependent GATA2 gain-of-function model to assess the impact of increased GATA2 dosage in LECs.

Shear stress signaling is a critical driver of lymphatic vascular development, and mutations in shear stress-sensing molecules, such as PIEZO1, are associated with human lymphedema ^1,23,26–35^. Because GATA2 and FOXC2 are both linked to mechanosensitive transcriptional programs, we reasoned that their dosage-sensitive interaction may converge on shear stress-responsive transcriptional programs ^15,30^. Hence, we performed our mechanistic studies in human LECs under static, oscillatory shear stress (OSS), and laminar shear stress (LSS) conditions. This approach enabled us to define how shear stress-dependent chromatin accessibility and gene expression programs are regulated by GATA2, FOXC2, and PROX1 in a human LEC context.

Together, these approaches reveal that GATA2 and FOXC2 cooperate in a dosage-sensitive manner to preserve a PROX1-centered shear stress-responsive transcriptional network required for lymphatic vascular development. These findings support a modifier model in which combined disruption of GATA2- and FOXC2-dependent regulatory inputs results in severe lymphatic vascular disease.

## RESULTS

### Gata2^+/-^ mice have fewer lymphatic- and lymphovenous valves

We generated embryonic day (E) 15.5 *Gata2^+/-^* embryos by breeding *Gata2^+/-^* and wild-type mice. We observed mild edema in 9 of 21 *Gata2^+/-^* embryos analyzed (**Figure 1A**). Edema was not observed in wild type embryos. Two pairs of LVVs are bilaterally located at the junction of jugular and subclavian veins, where they regulate lymph return to blood circulation. Defects in LVVs are frequently observed in mouse models of lymphedema ^22^. To assess whether LVVs are affected in *Gata2^+/-^* embryos, we collected E14.5, E15.5 and E16.5 embryos, cryo-embedded and sectioned them in frontal orientation, and performed immunohistochemistry using the LEC marker PROX1 and the pan-endothelial cell marker PECAM-1, both of which are robustly expressed in LVVs (**Figure 1B**, arrows). Among 23 wild-type embryos that were analyzed, 22 displayed 4 LVVs, while one embryo had 3 LVVs (**Figure 1C**). In contrast, approximately 40% of the 21 *Gata2^+/-^* embryos that were analyzed exhibited 2 or 3 LVVs. However, none of the *Gata2^+/-^* embryos had fewer than 2 LVVs.

**Figure 1:**
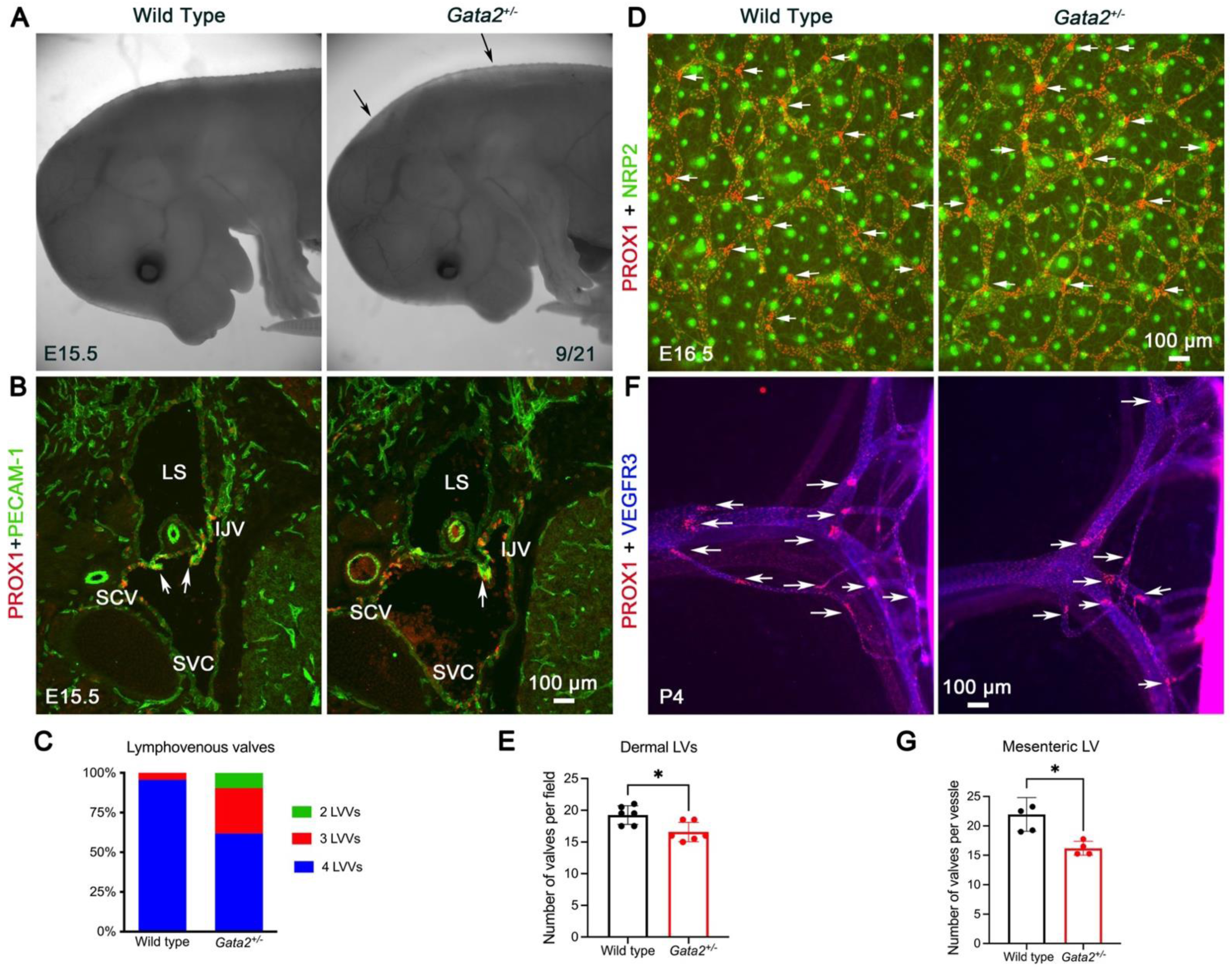
Lymphatic and lymphovenous valve development is affected by *Gata2*-heterozygosity. **(A)** Approximately 50% of E15.5 *Gata2^+/-^* embryos develops mild edema (arrows). **(B)** E15.5 wild-type and *Gata2^+/-^* embryos were frontally sectioned and immunohistochemistry was performed with the indicated antibodies. Two pairs of bilaterally located LVVs were normally observed in control embryos. One pair (arrows) is shown in the figure. In contrast, fewer LVVs were observed in a subset of *Gata2^+/-^*embryos. **(C)** E14.5, E15.5 and E16.5 wild-type and *Gata2^+/-^* embryos were analyzed and their LVVs were quantified. **(D)** Dorsal skin of E16.5 embryos was analyzed using the indicated antibodies. The PROX1^+^;NRP2^+^ structures are the lymphatic vessels, and the arrows indicate the LVs. The NRP2^+^ punctuated structures are sensory neurons. **(E)** Dermal LVs from E16.5 embryos were counted and quantified. **(F)** Mesenteric tissue from P4 pups was analyzed using the indicated antibodies. Arrows indicate the LVs. **(G)** Mesenteric LVs were counted and quantified. **Statistics**: (**C**) n=23 (n=9 E15.5, n= 14 E16.5) wild-type and n=21 *Gata2^+/-^* (n=3 E14.5, n=11 E15.5, n=7 E16.5) embryos were analyzed. (**E**) n=6 wild-type and n=6 *Gata2^+/-^* embryos were analyzed at E16.5. Two fields per skin sample were counted. Each dot represents the number of LVs per field. A Student’s unpaired *t*-test was performed for the statistical analysis. * p<0.05. (**G**) n=5 wild-type and n=4 *Gata2^+/-^* pups were analyzed. Three mesenteric vessels from each pup were analyzed. Each dot in the graph represents the average number of LVs per vessel. The Mann-Whitney test was performed for the statistical analysis. * p<0.05. Abbreviations: LS, lymph sacs; EJV, external jugular vein; SVC, superior vena cava; SCV, subclavian vein; IJV, internal jugular vein

We also examined the dermal lymphatic vessels in E16.5 *Gata2^+/-^* embryos and their control littermates. While no overt defects were observed in the overall organization of lymphatic vessels, the number of dermal LVs was reduced in *Gata2^+/-^* embryos (**Figure 1 D, E**). Similarly, fewer mesenteric LVs were observed in post-natal day (P) 4 *Gata2^+/-^* pups (**Figure 1F, G**). These findings indicated that *Gata2* haploinsufficiency is sufficient to reduce valve number, even in the absence of severe lymphatic patterning defects. This defect may underlie the mild edema observed in a subset of these animals.

### Lymphovenous and venous valves are absent in Gata2^+/-^;Foxc2^+/-^ mice

Reduced numbers of LVVs and LVs are a characteristic of *Foxc2^+/-^* mice ^22–24^. Chromatin immunoprecipitation using GATA2- or FOXC2-specific antibodies, followed by next-generation sequencing (ChIP-seq), has revealed that these TFs could co-occupy the regulatory elements of genes such as *PROX1* ^25^. Hence, we tested if the phenotype of *Gata2^+/-^* mice could be aggravated in the *Foxc2*-heterozygous background. We bred *Gata2^+/-^* to *Foxc2^+/-^* mice, and 100% of the E15.5 *Gata2^+/-^;Foxc2^+/-^* embryos generated by this cross had edema (**Figure 2A**). Analysis of jugular and subclavian vein junction of E14.5, E15.5 and E16.5 embryos revealed the absence of LVVs in double heterozygous embryos (**Figure 2B-D**). Despite the lack of LVVs and the presence of dilated lymph sacs, no blood was observed within the lymph sacs of *Gata2^+/-^;Foxc2^+/-^* embryos (**Figure 2B**), indicating that the absence of LVVs does not result in a blood-filled lymphatic phenotype.

**Figure 2:**
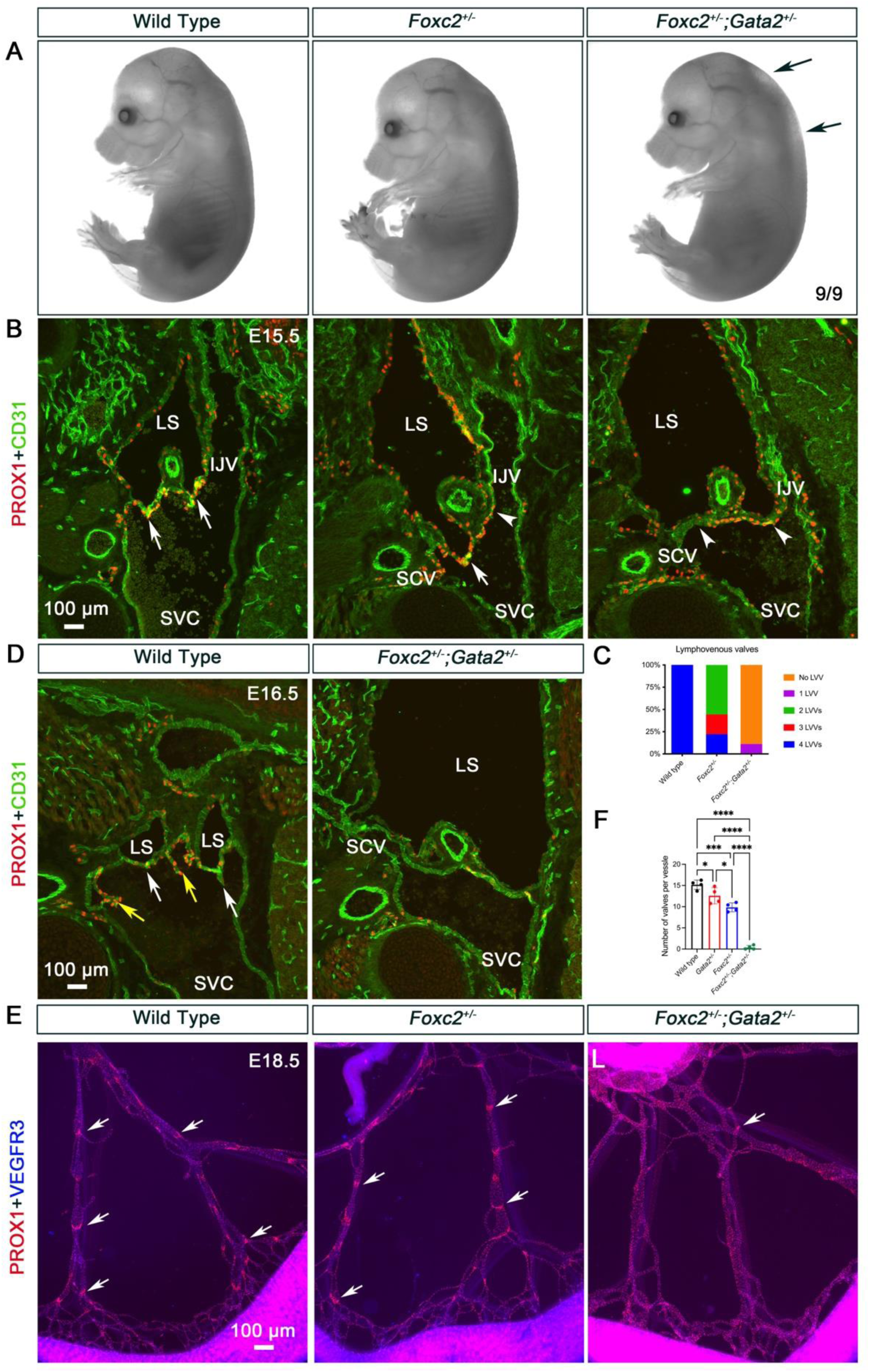
Lymphatic, venous and lymphovenous valves are absent in *Gata2^+/-^;Foxc2^+/-^* embryos. *Foxc2^+/-^* and *Gata2^+/-^* mice were bred and their embryos were analyzed for valves at various developmental time points. **(A)** Edema was observed in 100% of E15.5 *Gata2^+/-^*;*Foxc2^+/-^* embryos, although it was less frequently observed in single heterozygous embryos. **(B)** E15.5 embryos were frontally sectioned to observe the bilaterally located LVVs. One side with a pair of LVVs (arrows) is shown. While control embryos had the normal number of LVVs, fewer LVVs were observed in *Foxc2^+/-^* embryos. In contrast, no LVVs were observed in *Gata2^+/-^*;*Foxc2^+/-^* embryos. Arrowheads indicate the locations where LVVs should have formed. **(C)** LVVs in E15.5 embryos were counted and quantified. **(D)** E16.5 embryos were frontally sectioned to identify LVVs (white arrows) and venous valves (yellow arrows). LVVs and venous valves were absent in *Gata2^+/-^*;*Foxc2^+/-^* embryos and their lymph sacs were massively dilated. **(E)** Mesenteric lymphatic vessels of E18.5 embryos were analyzed. LVs (arrows) were observed in control, *Foxc2^+/-^* and *Gata2^+/-^* embryos, but only rarely in *Gata2^+/-^*;*Foxc2^+/-^*embryos. **Statistics**: (**A-C**) n=9 wild-type, n=9 *Foxc2^+/-^* and n=9 *Gata2^+/-^;Foxc2^+/-^* embryos at E15.5 were analyzed. (**D**) n= 14 wild-type and n=4 *Gata2^+/-^;Foxc2^+/-^* embryos at E16.5 were analyzed. (**E, F**) n=4 wild-type, n=4 *Foxc2^+/-^*, n=4 *Gata2^+/-^* and n= 5 *Gata2^+/-^;Foxc2^+/-^* embryos at E18.5 were analyzed. A one-way ANOVA was performed for statistical analysis. * p<0.05, ***p<0.005, **** p<0.001. Abbreviations: LS, lymph sacs; EJV, external jugular vein; SVC, superior vena cava; SCV, subclavian vein; IJV, internal jugular vein

Venous valves resemble LVVs at the molecular level ^22,36–38^. They develop in the vicinity of LVVs at the junctions of the internal jugular vein, external jugular vein and subclavian vein with the superior vena cava. PROX1^high^ venous valve progenitors can be seen in this location in control and *Gata2^+/-^;Foxc2^+/-^* embryos at E15.5 (**Figure 2B**, arrowheads). The venous valve progenitors developed into venous valves in control embryos at E16.5 (**Figure 2D**, yellow arrows). In contrast, neither venous valves nor PROX1^high^ cells were observed in E16.5 *Gata2^+/-^;Foxc2^+/-^* embryos (**Figure 2D**). This observation indicates that the specification of valvular endothelial cells is not affected in *Gata2^+/-^;Foxc2^+/-^* embryos, although their subsequent survival and morphogenesis are compromised.

### Lymphatic vessel maturation is defective, and lymphatic valves are reduced in Gata2^+/-^;Foxc2^+/-^ mice

LECs originate primarily from embryonic veins during the E9.5-12.5 developmental window ^39^. These cells give rise to the lymph sacs and subsequently migrate to various organs, where they assemble into the primitive lymphatic plexus. This plexus then undergoes stepwise maturation into lymphatic capillaries, pre-collecting vessels, and collecting lymphatic vessels. Proper development and maturation of these vessels are prerequisites for the formation of LVs, which are located primarily within pre-collecting and collecting lymphatic vessels. FOXC2 and GATA2 are essential for these processes, as mice deficient in either one of these TFs display immature, dilated lymphatic vessels with aberrant recruitment of vascular smooth muscle cells ^14–16,21,40^.

The mesenteric lymphatic vessels of E18.5 *Gata2^+/-^;Foxc2^+/-^* embryos had a mesh-like architecture (**Figure 2E**). While LVs were reduced in numbers in *Gata2^+/-^* and *Foxc2^+/-^*embryos, they were nearly absent in *Gata2^+/-^;Foxc2^+/-^* embryos (**Figure 2E, F**). Only a few PROX1^high^ cells were observed (**Figure 2E**, arrow).

Compared to littermates, *Gata2^+/-^;Foxc2^+/-^* embryos remained edematous at E18.5 (**Figure 3A**, arrow). Analysis of the skin revealed lymphatic vessels with bulbous architecture in *Gata2^+/-^;Foxc2^+/-^* embryos (**Figure 3B**). These vessels were abnormally associated with α-smooth muscle actin^+^ (α-SMA^+^) vascular smooth muscle cells (**Figure 3C**, arrows). Overall, the dermal and mesenteric lymphatic vessel phenotypes of *Gata2^+/-^;Foxc2^+/-^* mice suggest a defect in the maturation of these structures. These phenotypes are highly reminiscent of those reported in *Foxc2^-/-^* embryos ^40^.

**Figure 3:**
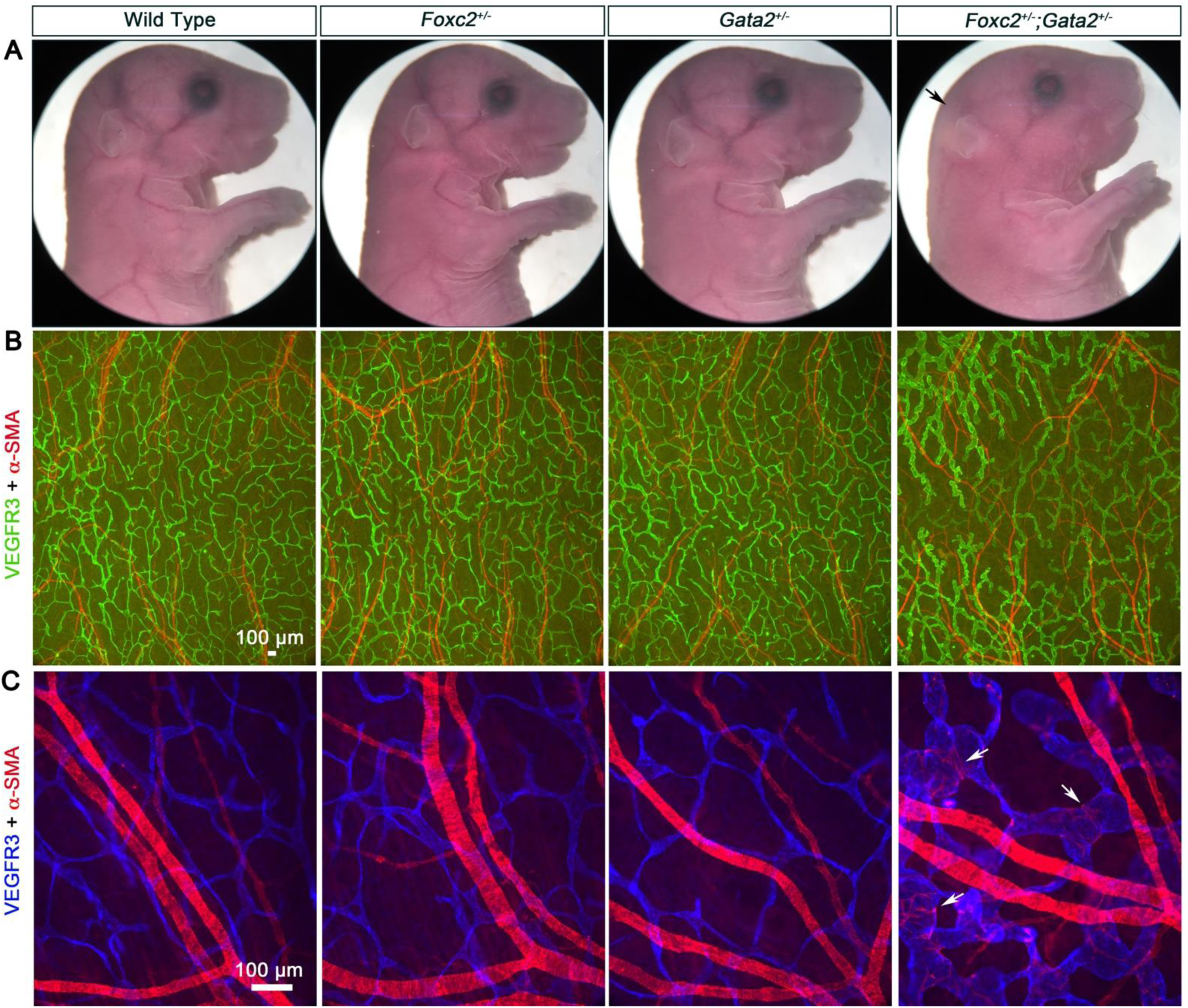
Lymphatic vessels of *Gata2^+/-^;Foxc2^+/-^* embryos are dysplastic. **(A)** Edema was observed in E18.5 *Foxc2^+/-^*;*Gata2^+/-^* embryos (arrow), although it was not obvious in single heterozygous embryos at this stage. **(B)** Dorsal skin of E18.5 *Foxc2^+/-^*;*Gata2^+/-^* embryos was analyzed using the indicated antibodies. The lymphatic vessels of double heterozygous embryos were abnormally shaped with a “bulbous” architecture. **(C)** At higher magnification, abnormal recruitment of α-smooth muscle actin^+^ (α-SMA) vascular smooth muscle cells was observed on the dermal lymphatic vessels of E18.5 *Foxc2^+/-^*;*Gata2^+/-^* embryos. **Statistics**: n=4 wild-type, n=4 *Foxc2^+/-^*, n=4 *Gata2^+/-^* and n= 5 *Gata2^+/-^;Foxc2^+/-^* embryos were analyzed.

### The lymphatic valves of adult Gata2^+/-^;Foxc2^+/-^ mice are dysfunctional

We generated postnatal mice by breeding *Gata2^+/-^* and *Foxc2^+/-^* mice. From 28 litters, we obtained 164 live pups at postnatal day (P)10, of which only 10 were *Gata2^+/^;Foxc2^+/-^*, significantly fewer than the 41 that were expected (**Figure 4A**). In contrast, *Foxc2^+/-^* and *Gata2^+/-^* mice were each obtained close to the expected frequency. We observed several dead *Gata2^+/^;Foxc2^+/-^* pups in the cage immediately after birth. Hence, the survival of *Gata2^+/^;Foxc2^+/-^* mice is strikingly reduced due to perinatal lethality.

**Figure 4.**
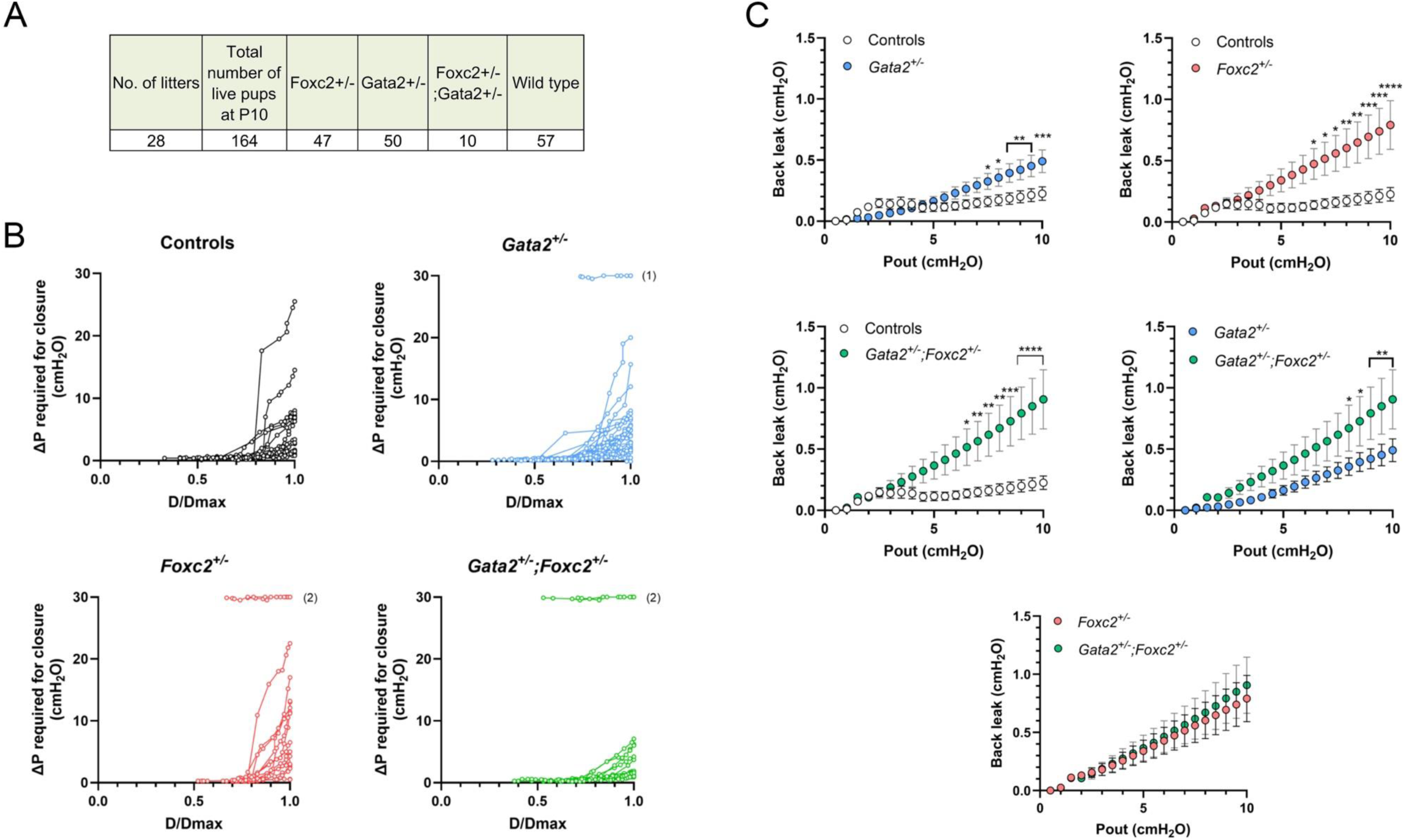
Lymphatic valves of *Gata2^+/-^*, *Foxc2^+/-^* and *Gata2^+/-^;Foxc2^+/-^* mice are defective. A) Survival table. *Gata2^+/-^* and *Foxc2^+/-^* mice were bred and the number of surviving pups at P10 was quantified. *Gata2^+/-^* ;*Foxc2^+/-^* pups were recovered at a frequency below Mendelian expectations (41 expected versus 10 observed). Forty-one pups were expected for each genotype. **B) Closure tests.** Closure tests for control valves (WT littermates of the other three genotypes), showing values of ΔP plotted against the initial (normalized) vessel diameter (D/Dmax) at different levels of Pin. ΔP was calculated from Pout-Pin at the moment during the Pout ramp that the valve closed. The sets of ΔP values obtained for each valve are connected by a solid line. All but two control valves closed at ΔP < 8 cmH2O, even at the maximal diameters, the other two required a higher ΔP but closed. One *Gata2^+/-^* valve, two *Foxc2^+/-^* valves and two *Gata2^+/-^;Foxc2^+/-^* valves never closed even at ΔP = 30 cmH_2_O. **C) Back leak tests.** Comparison of back leak (Psn-Pin) as a function of Pout for various genotypes of mice. *Gata2^+/-^;Foxc2^+/-^* valves were significantly leakier than *Gata2^+/-^*, but not *Foxc2^+/-^* valves. **Statistics: (A)** Chi-square test revealed a highly significant reduction in survival of double heterozygous pups with χ^2^=31.25, degrees of freedom=1 and p=2.3 X 10^-8^, significantly below Mendelian expectation. **(B)** n=8 mice, n=18 valves for controls; n=12 mice, n=29 valves for *Gata2^+/-^*; n=4 mice, n=15 valves for *Foxc2^+/-^*; n=5 mice, n=15 valves for *Gata2^+/-^ ;Foxc2^+/-^.* **(C)** Comparisons were made using 2-way ANOVAs with Tukey’s post-hoc tests. * = p <0.05; ** = p<0.01; *** = p<0.001; **** = P <0.0001. n=8 mice, n=18 valves for controls; n=12 mice, n=29 valves for *Gata2^+/-^*; n=4 mice, n=15 valves for *Foxc2^+/-^*; n=5 mice, n=15 valves for *Gata2^+/-^;Foxc2^+/-^*.

Although most *Gata2^+/-^;Foxc2^+/-^* mice die perinatally, a small fraction survives into adulthood, likely due to less severe defects. We studied the structure and function of LVs from the surviving *Gata2^+/^;Foxc2^+/-^* mice.

#### Functional tests on popliteal lymphatic valves

Popliteal lymphatic vessels from control, *Gata2^+/-^*, *Foxc2^+/-^* and the surviving *Gata2^+/^;Foxc2^+/-^* mice were dissected and cleaned for ex vivo quantification of LV function. A lymphatic vessel containing a single valve was cannulated at each end for independent control of inflow pressure (Pin) and outflow pressure (Pout), as shown in **Supplementary Figure 1A**. Tests were conducted in Ca^2+^-free Krebs solution to eliminate pressure spikes from spontaneous contractions. Two related protocols were performed to determine: 1) the transvalvular pressure gradient (ΔP) required to close a valve (a measure of leaflet stiffness and valve competency); 2) the back leak pressure across a closed valve (a measure of how well the closed valve sealed). The protocols are illustrated by the images and recordings in **Supplementary Figure 1**.

Closure tests on popliteal lymphatic valves of *Gata2^+/-^, Foxc2^+/-^*, and *Gata2^+/-^;Foxc2^+/-^*<u>mice:</u> Because the ΔP for closure varied with the initial diameter/pressure of the vessel, a complete “closure curve” was determined by setting Pin at 10 different levels between 0.1 and 10 cmH_2_O and raising Pout ramp-wise each time until the valve closed; the adverse ΔP (i.e., Pout-Pin) required for closure at each level of Pin was then plotted as a function of the normalized diameter; the curve describing this relationship is shown for 18 control valves (WT littermates for the various genotypes) in **Figure 4B**. Two of 18 vessels required higher than normal pressures for valve closure at the maximum diameter, but still closed, which is consistent with our previous studies ^41,42^. In contrast, 1 of 29 *Gata2^+/-^*, 2 of 15 *Foxc2^+/-^*, and 2 of 15 *Gata2^+/-^;Foxc2^+/-^* valves never closed even at ΔP = 30 cmH_2_O (**Figure 4B**), the maximum level allowed by the transducers in our servo-control system, reflecting the behavior of valves that were completely incompetent.

Back leak measurements on popliteal lymphatic valves of *Gata2^+/-^, Foxc2^+/-^*, and *Gata2^+/-^;Foxc2^+/-^* mice: Back leak across a closed LV was measured in the same vessels. Starting with Pin=Pout=0.5 cmH_2_O, Pout was raised ramp-wise to 10 cmH_2_O and if the valve did not close before ΔP = 1 cmH_2_O, the outflow line was gently tapped to produce pressure spikes and encourage valve closure. This procedure was repeated 3 times for each valve while recording Psn upstream from the closed valve. For data analysis, the continuous recordings of Psn values were binned in 0.5 cmH_2_O intervals of Pout and the sets of binned data for different valves were compared either to the control group or to another genotype using a 2-way ANOVA. The datasets for back leak (calculated from Psn-Pin) as a function of Pout were then plotted in **Figure 4C**, with significance between groups at each pressure interval determined using Tukey’s multiple-comparison tests.

*Gata2^+/-^* valves exhibited a modest but significant degree of back leak once Pout exceeded 7 cmH_2_O (**Figure 4C**). *Foxc2^+/-^* valves showed more significant back leak once Pout exceeded 6 cmH_2_O. We observed more severe back leak in popliteal LVs of *Foxc2^+/-^*mice compared to previous studies that were performed on mesenteric LVs of these mice ^23,43^. This observation suggests that regional differences in LV function occur and this distinction may help explain the peripheral lymphedema, but not chylous ascites, that is observed in lymphedema distichiasis patients. We recently reported similar regional differences among LVs in the mesentery, specifically between those located in jejunum-draining versus ileum-draining lymphatic vessels in mice lacking S1PR1 in LECs ^44^.

*Gata2^+/-^;Foxc2^+/-^* valves also showed significant back leak once Pout exceeded 6 cmH_2_O. Back leak across *Gata2^+/-^;Foxc2^+/-^* valves was more severe than for *Gata2^+/-^*valves and was significant at the 5 highest Pout levels. There were no significant differences in back leak between *Foxc2^+/-^* valves and *Gata2^+/-^;Foxc2^+/-^* valves. Thus, the valves of surviving *Gata2^+/-^;Foxc2^+/-^* mice were functionally similar to those of *Foxc2^+/-^*mice. These findings further suggest that the more severely affected *Gata2^+/-^;Foxc2^+/-^*mice may have died prior to analysis.

#### Leaflet length analysis

Finally, we analyzed leaflet dimensions across individual valves of each genotype. Leaflet lengths were measured from high-magnification brightfield images acquired at the top and bottom surfaces of each vessel, according to the structures identified and defined in a recent publication ^41^. Representative images from each surface are shown in **Figure 5A** for wild-type, *Gata2^+/-^, Foxc2^+/-^* and *Gata2^+/-^ ;Foxc2^+/-^* valves. As quantified in **Figure 5B**, wild-type control valves had average leaflet lengths of ∼110 μm. By contrast, *Foxc2^+/-^* and *Gata2^+/-^;Foxc2^+/-^* valves had significantly shorter average leaflet lengths than controls.

**Figure 5.**
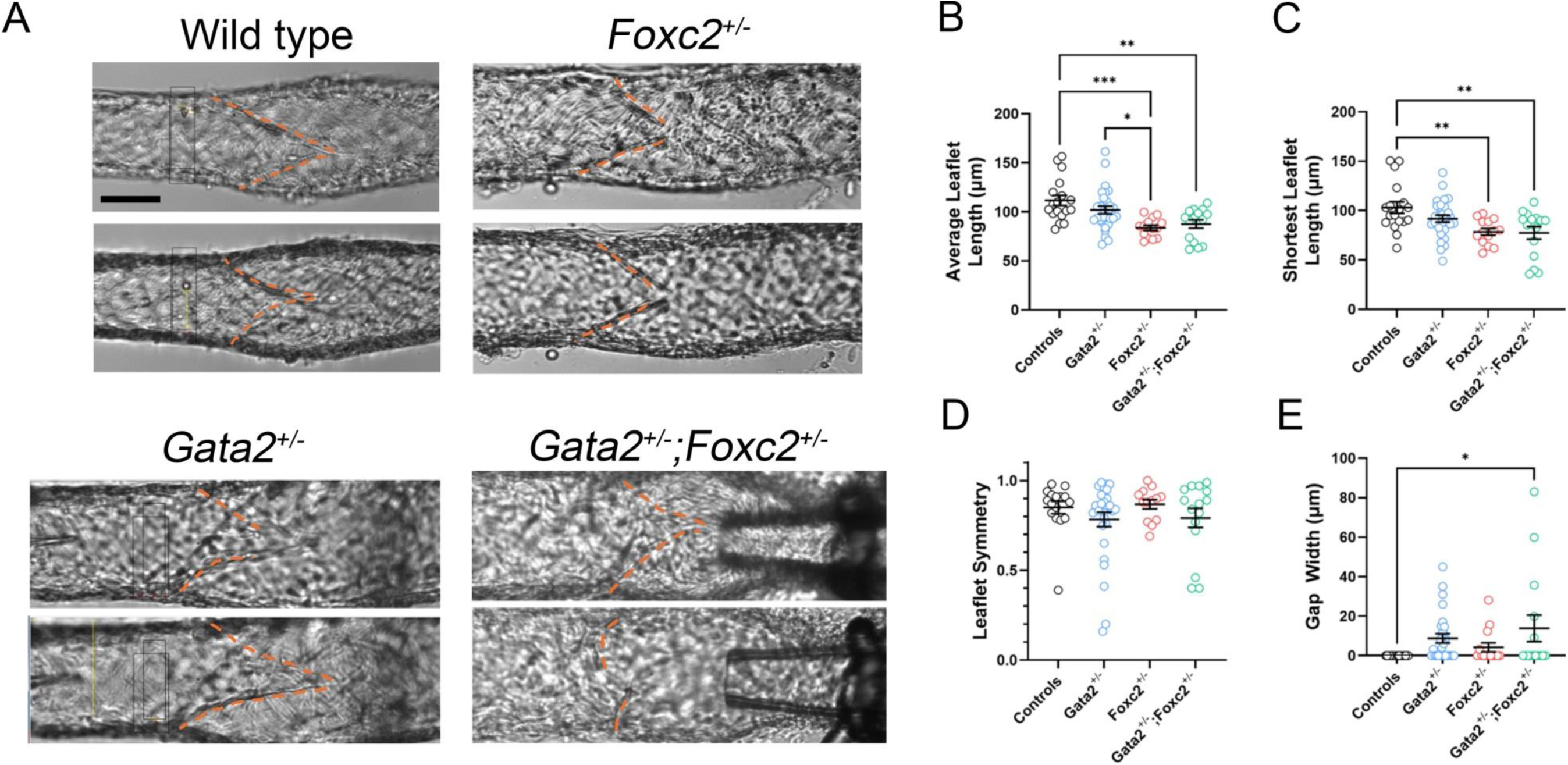
Lymphatic valve leaflets of *Gata2^+/-^, Foxc2^+/-^* and *Gata2^+/-^;Foxc2^+/-^* mice are shorter. **A**) Images of the upper and lower surfaces of valves from mice of the various genotypes, showing the insertion points of the leaflets at the two respective focal planes. Dotted orange lines were added (and offset by a few pixels) to trace the leaflets. Calibration bar = 50 μm applies to all images. **B**) Analysis of average leaflet length (average of the two leaflets in each valve) by genotype. **C**) Analysis of the shortest leaflet length (the shorter of the two leaflets in each valve) by genotype. **D**) Analysis of length symmetry by genotype. **E**) Analysis of gap width (the shorter of gap width for each valve) by genotype. **Statistics: (B-E)** Each dot in the graphs indicates an individual valve. Comparisons were made using a 1-way ANOVA with Tukey’s post-hoc tests. * = p <0.05; ** = p<0.01; *** = p<0.001. Unmarked comparisons were n.s.

We next plotted the lengths of the shorter leaflets from each valve (**Figure 5C**), reasoning that even a single short leaflet could limit valve closure and/or sealing. By this metric, only *Foxc2^+/-^* and *Gata2^+/-^;Foxc2^+/-^* valves were significantly shorter than wild-type controls. There were no significant differences in leaflet symmetry (**Figure 5D**). We also measured the gap width between the ends of the leaflets because gap width correlated strongly with the degree of back leak in mice with connexin deficiencies ^41^. In our analysis, gap width was increased not only in valves with one or both shortened leaflets, but also in many non-wild-type valves with apparently normal leaflet lengths. These observations support the concept that valve competency depends not only on leaflet length, but also on leaflet apposition and maturation of valve geometry. Consistent with this interpretation, gap width was significantly increased only in *Gata2^+/-^;Foxc2^+/-^* valves (**Figure 5E**).

Taken together, these findings reveal that the LVs of *Gata2^+/-^, Foxc2^+/-^* and *Gata2^+/-^ ;Foxc2^+/-^* mice exhibit varying degrees of structural abnormalities, such as shortened leaflets and gaps between the leaflet tips, that compromise their ability to close and prevent back leak.

### GATA2 overexpression disrupts PROX1-dependent lymphatic vascular development

The phenotypes of *Gata2^+/-^;Foxc2^+/-^* mice, including the absence of LVVs, LVs, and venous valves, as well as reduced viability, closely resemble those of *Prox1^+/-^* mice ^22,36,45^. In fact, a recent ChIP-seq analysis identified a conserved regulatory element in the *PROX1* locus that is co-occupied by PROX1, FOXC2, GATA2 and NFATC1 ^25^. Surprisingly, however, PROX1 expression is paradoxically elevated in the lymphatic vessels of *Gata2^+/-^;Foxc2^+/-^* mice (**Figure 2E**). PROX1 expression is also elevated in mice lacking GATA2 or FOXC2 in LECs ^15,16,21,32,40^. Consistent with this, FOXC2 and GATA2 expression decreases, whereas PROX1 expression increases, as LEC progenitors differentiate and migrate out from embryonic cardinal vein ^22^. Collectively, these findings point to a more complex regulatory relationship between PROX1, FOXC2 and GATA2.

To further dissect the interplay between GATA2 and PROX1, we generated a Cre-dependent GATA2 gain-of-function mouse model (Gata2^GOF^; **Figure 6A**) and crossed it with *Prox1^+/GFPCre^* mice to drive GATA2 overexpression specifically in PROX1^+^ cells, including the developing LECs ^46^. *Prox1^+/GFPCre^*;Gata2^GOF^ embryos were collected at E11.5 and E14.5 for phenotypic analysis (**Figure 6B**).

**Figure 6:**
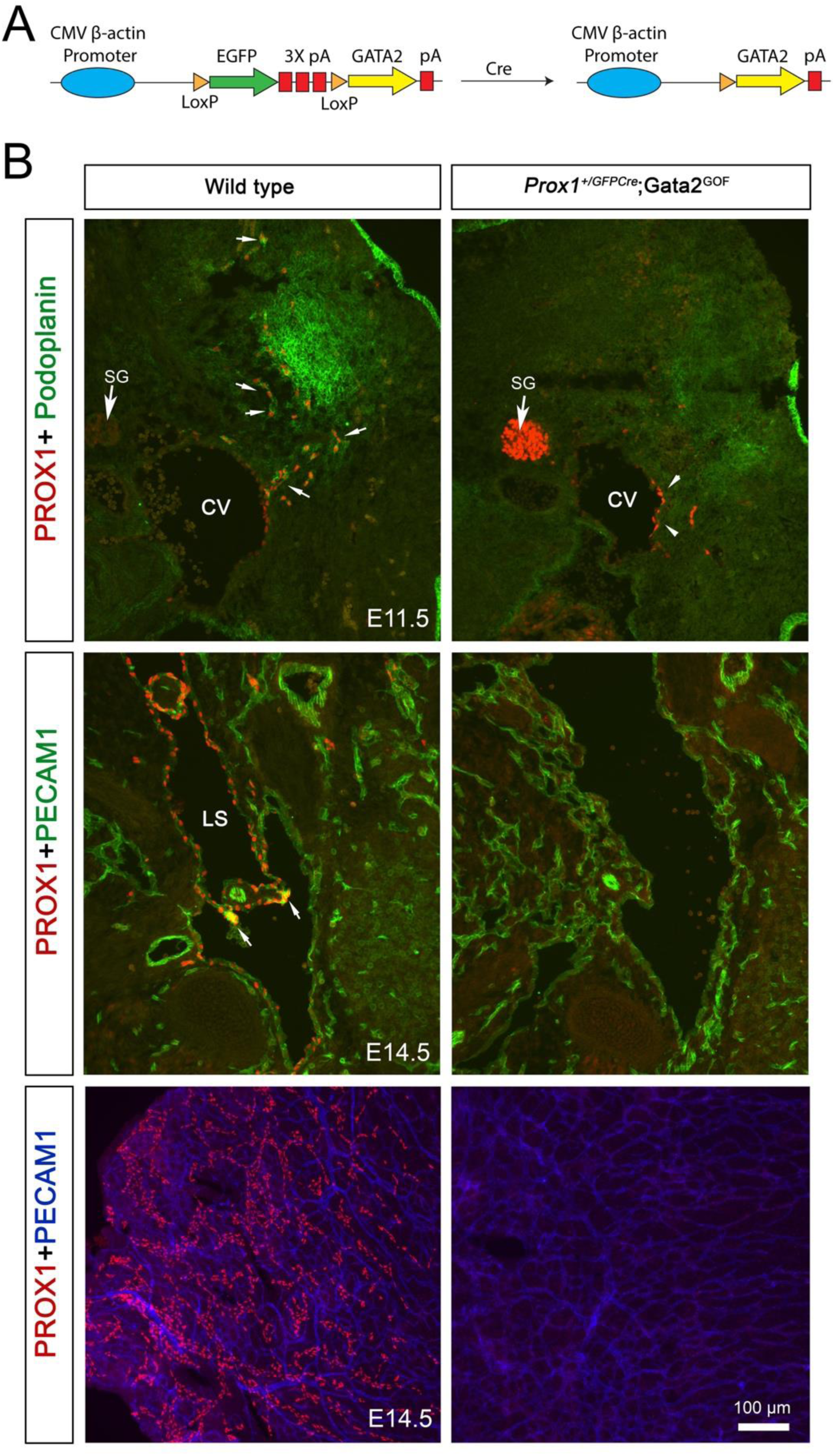
GATA2 constrains PROX1 expression in vivo. (A) Design of the Cre-dependent GATA2 overexpression model (Gata2^GOF^). EGFP is constitutively expressed in the absence of Cre recombinase. Upon Cre-mediated deletion of the EGFP cassette, GATA2 is expressed. (B) Overexpression of GATA2 constrains PROX1 expression in LECs. *Prox1^+/GFPCre^*;Gata2^GOF^ embryos and wild-type littermates were analyzed at E11.5 or E14.5. At E11.5, PROX1^+^ LEC progenitors were seen on the cardinal vein (CV), and PROX1^+^ podoplanin^+^ LECs were seen migrating from the CV of wild-type embryos. In contrast, few LEC progenitors and LECs were observed in *Prox1^+/GFPCre^*;Gata2^GOF^ embryos (arrowheads). We had to substantially increase the camera exposure to observe these PROX1^+^ cells. Consequently, PROX1 expression in the sympathetic ganglia (SG) was dramatically increased. At E14.5, lymph sacs (LS) and lymphovenous valves (arrows) were seen in wild-type embryos but were completely absent in *Prox1^+/GFPCre^*;Gata2^GOF^ embryos. Consistent with this observation, while lymphatic vessels were observed in the dorsal skin of control embryos, none were seen in *Prox1^+/GFPCre^*;Gata2^GOF^ embryos. **Statistics:** (B) Images are representative of n=4 samples per genotype per stage

At E11.5, the PROX1^+^ LEC progenitors were readily identified in the cardinal vein, and PROX1^+^ podoplanin^+^ LECs were seen migrating away from the veins of wild-type embryos, as previously described (**Figure 6B**, arrows) ^22,47^. In striking contrast, *Prox1^+/GFPCre^*;Gata2^GOF^ littermates exhibited a marked downregulation of both PROX1 and podoplanin expression in the sparse LEC progenitors and LECs that were detected. By E14.5, lymph sacs and LVVs were present in wild-type embryos but were entirely absent in *Prox1^+/GFPCre^*;Gata2^GOF^ littermates. Accordingly, lymphatic vessels visible in the dorsal skin of wild-type embryos were also absent in the mutants (**Figure 6B**).

We previously showed that FOXC2 overexpression in LECs disrupts LV formation, resulting in a phenotype resembling *Prox1^+/-^* mice ^32^. Together, these results demonstrate that GATA2 and FOXC2 overexpression impairs lymphatic vascular development and are associated with reduction of PROX1, supporting a model in which GATA2 and FOXC2 dosage constrains PROX1-dependent developmental programs within a functional range.

### Shear stress defines distinct chromatin accessibility landscapes and engages context-dependent transcriptional networks

Having established that GATA2 and FOXC2 cooperate to regulate lymphatic vascular development, at least in part through PROX1, we next sought to identify the downstream transcriptional programs regulated by these factors. Shear stress is a key regulator of vascular biology. In the lymphatic vasculature, LSS induces KLF2 and KLF4 expression, suppresses Notch signaling, promotes LEC quiescence, and enhances responsiveness to VEGF-C ^34,35,48^. GATA2 and FOXC2 are mechanosensitive TFs whose expression is enhanced by OSS ^1,15,30–32,49^. GATA2 regulates the OSS-dependent FOXC2 expression, whereas FOXC2 promotes lymphatic vascular integrity and quiescence.

To define the chromatin landscape underlying these responses, we performed ATAC-seq on primary human LECs (HLECs) cultured under static, LSS, or OSS conditions. Principal component analysis (PCA) of accessible genomic regions revealed clear segregation among the three conditions, along with tight clustering of biological replicates, indicating high reproducibility and demonstrating that shear stress is a major determinant of chromatin accessibility (**Figure 7A**).

**Figure 7:**
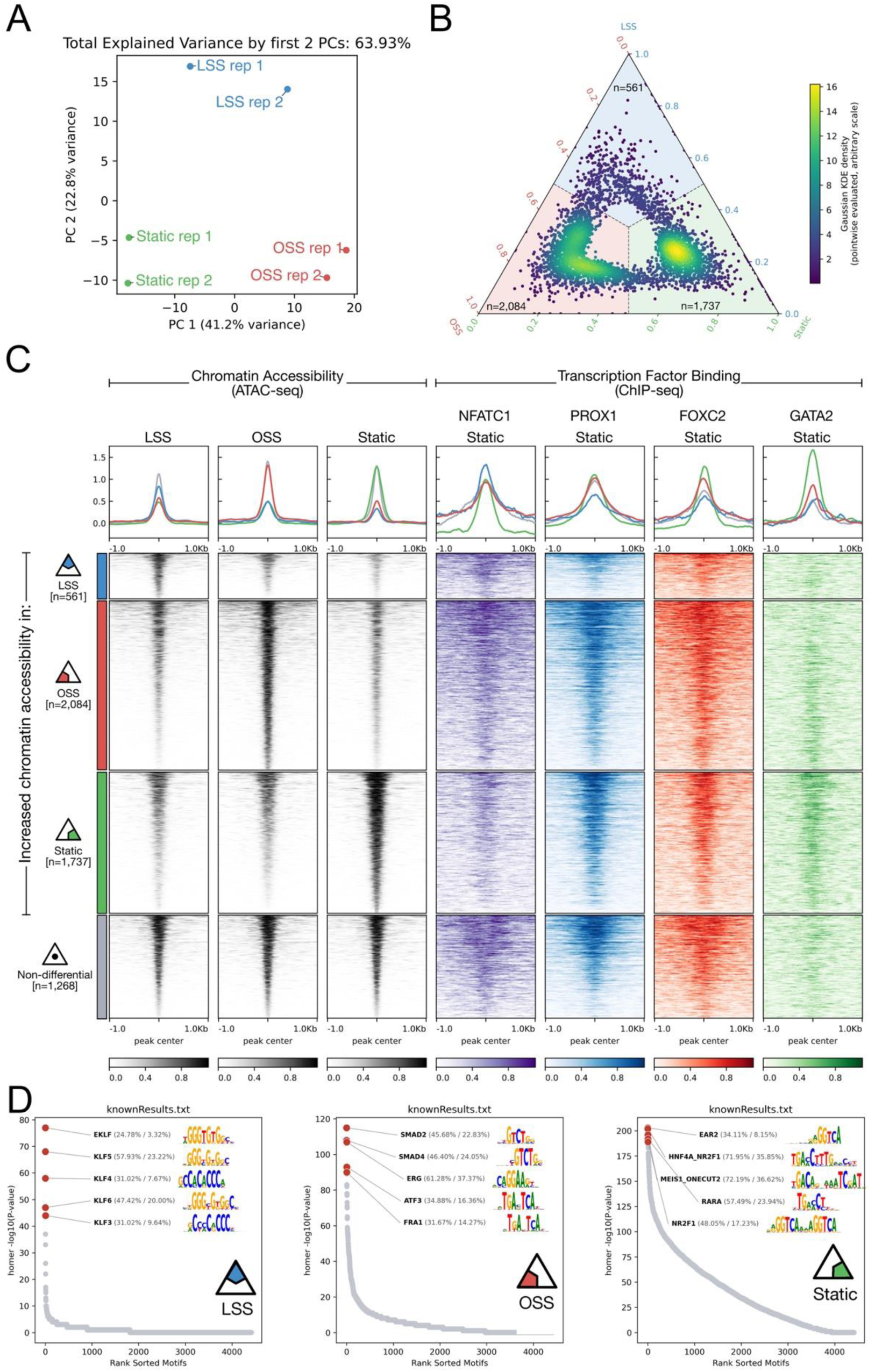
Shear stress induces distinct chromatin accessibility landscapes with selective transcription factor engagement and functional output. (A) Principal component analysis (PCA) of ATAC-seq datasets from HLECs cultured under static, LSS or OSS conditions. PCA was performed on the 3,000 most variable open chromatin regions, ranked based on standard deviation across all replicates and conditions, and using normalized read counts across all open chromatin regions (intervals). Each point represents an individual biological replicate, colored by condition (red, static; green, LSS; blue, OSS). PC1 and PC2 account for the indicated percentages of the total variance. Replicates cluster tightly within condition, demonstrating high reproducibility. Clear segregation of static, LSS and OSS samples indicates distinct shear stress conditions induce condition-specific chromatin accessibility profiles. (B) Ternary plot comparing ATAC-seq signal intensities across static, OSS and LSS conditions. The plot is partitioned based on relative enrichment across the three states: 561 regions (top vertex) are enriched under LSS, 2,084 regions (left vertex) are enriched under OSS, and 1,737 regions (right vertex) are enriched under static condition, highlighting condition-specific chromatin accessibility landscapes. (C) Peak-centered signal enrichment heatmap (tornado plots) depicting shear condition-specific and shear-independent chromatin accessibility sites identified by ATAC-seq, alongside ChIP-seq signals for NFATC1, PROX1, FOXC2 and GATA2 profiled under static conditions. Regions are anchored at ATAC-seq peak centers and ranked by signal intensity. These analyses show that all four TFs broadly co-occupy accessible chromatin irrespective of shear condition. Notably, GATA2 exhibits preferential enrichment at sites overrepresented under static condition. (D) TF binding motifs enriched within shear condition-specific chromatin accessibility sites. Motif analysis revealed KLF family motifs in LSS-associated regions when compared to non-differential sites. SMAD, ERG, and FRA1 motifs were enriched in OSS-associated regions, whereas nuclear hormone receptor motifs were enriched in static condition. **Statistics:** n=2 replicates of ATAC-seq were performed for each of the three (static, LSS or OSS) conditions.

To determine whether these shear stress-dependent changes in chromatin accessibility were associated with corresponding transcriptional changes, we performed bulk RNA-seq on HLECs cultured under static, LSS, and OSS conditions. Volcano plots comparing static HLECs with either LSS- or OSS-exposed HLECs revealed condition-specific changes in chromatin accessibility, identifying genomic intervals most strongly increased or decreased in each condition (**Supplementary Figure 2A**). Genomic intervals significantly enriched under specific shear stress conditions are listed in **Supplementary Tables 1, 2**.

We next identified differentially expressed genes in HLECs cultured under static versus LSS and static versus OSS conditions (**Supplementary Tables 3, 4**) and integrated this data with the ATAC-seq results using Gene Set Enrichment Analysis (GSEA). This analysis revealed significant, directionally consistent enrichment: genes associated with increased chromatin accessibility were enriched among upregulated transcripts, whereas genes associated with decreased accessibility were enriched among downregulated transcripts (**Supplementary Figure 2B**). Thus, shear stress-induced chromatin remodeling is globally aligned with gene expression changes in HLECs, consistent with a regulatory relationship between chromatin state and transcriptional output.

We generated a ternary plot to simultaneously compare chromatin accessibility across static, LSS, and OSS conditions based on per-interval mean signal proportions (**Figure 7B**). Each interval was assigned to one of three regions defined by the dominant compositional component (i.e., the condition with the largest fraction of total signal per interval). This partitioning yielded 561 intervals assigned to LSS, 2,084 intervals to OSS, and 1,737 intervals assigned to static condition (**Figure 7B** and **Supplementary Tables 5-7**). These condition-specific intervals were intersected with published ChIP-seq datasets for PROX1, FOXC2, NFATC1, and GATA2 in HLECs under the static conditions ^25^. All four TFs were generally associated with accessible chromatin irrespective of shear stress condition, consistent with their roles as core regulators of LEC identity (**Figure 7C**).

To identify additional TFs involved in shear stress-responsive gene regulation, accessible chromatin intervals from each condition were subjected to unbiased motif enrichment analysis (**Figure 7D** and **Supplementary Tables 8-11**). KLF motifs were dominant among the top-enriched motifs at sites enhanced by LSS. In contrast, SMAD, ERG (ETS), and FRA1 (AP-1) motifs were enriched in OSS-associated regions, whereas nuclear hormone receptor motifs were enriched under static conditions. The enrichment of ERG/ETS motifs in OSS-associated regions is consistent with recent evidence that ETS factors cooperate with valve-enriched TFs to regulate LV gene programs ^50,51^.

GATA2 can co-occupy AP-1 binding sites, while FOXC2 functions with ETS factors at shared regulatory elements ^52,53^. Similarly, PROX1 interacts with nuclear hormone receptors, ETS factors, and KLF2 to modulate transcriptional outputs ^34,35,46,54–56^. Collectively, these data support a model in which PROX1, GATA2, and FOXC2 occupy broadly accessible chromatin, while shear stress selectively engages distinct co-regulatory networks that drive condition-specific transcriptional outputs.

### PROX1 and FOXC2 differentially regulate shear stress-responsive genes under OSS

Having found that shear stress induces distinct chromatin accessibility landscapes while PROX1, FOXC2, and GATA2 remain broadly associated with accessible chromatin, we next asked how these core TFs regulate shear-responsive gene expression. We focused on their relationship with KLF2 and KLF4, two key TFs implicated in shear stress-responsive gene expression program.

RNA-seq analysis revealed that both LSS and OSS can enhance *KLF2* and *KLF4* expression in HLECs (**Supplementary Tables 3, 4)**. In fact, KLF motifs were enriched in ATAC-seq datasets under both LSS and OSS conditions when compared to static conditions (**Supplementary Figure 3 and Supplementary Tables 8, 10**). Analysis of the *KLF2* locus revealed that both LSS and OSS markedly increased chromatin accessibility across the gene body (**Figure 8A**, boxed region). Motif analysis of this region identified numerous KLF-binding sites, consistent with potential transcriptional autoregulation (**Supplementary Figure 4**).

**Figure 8:**
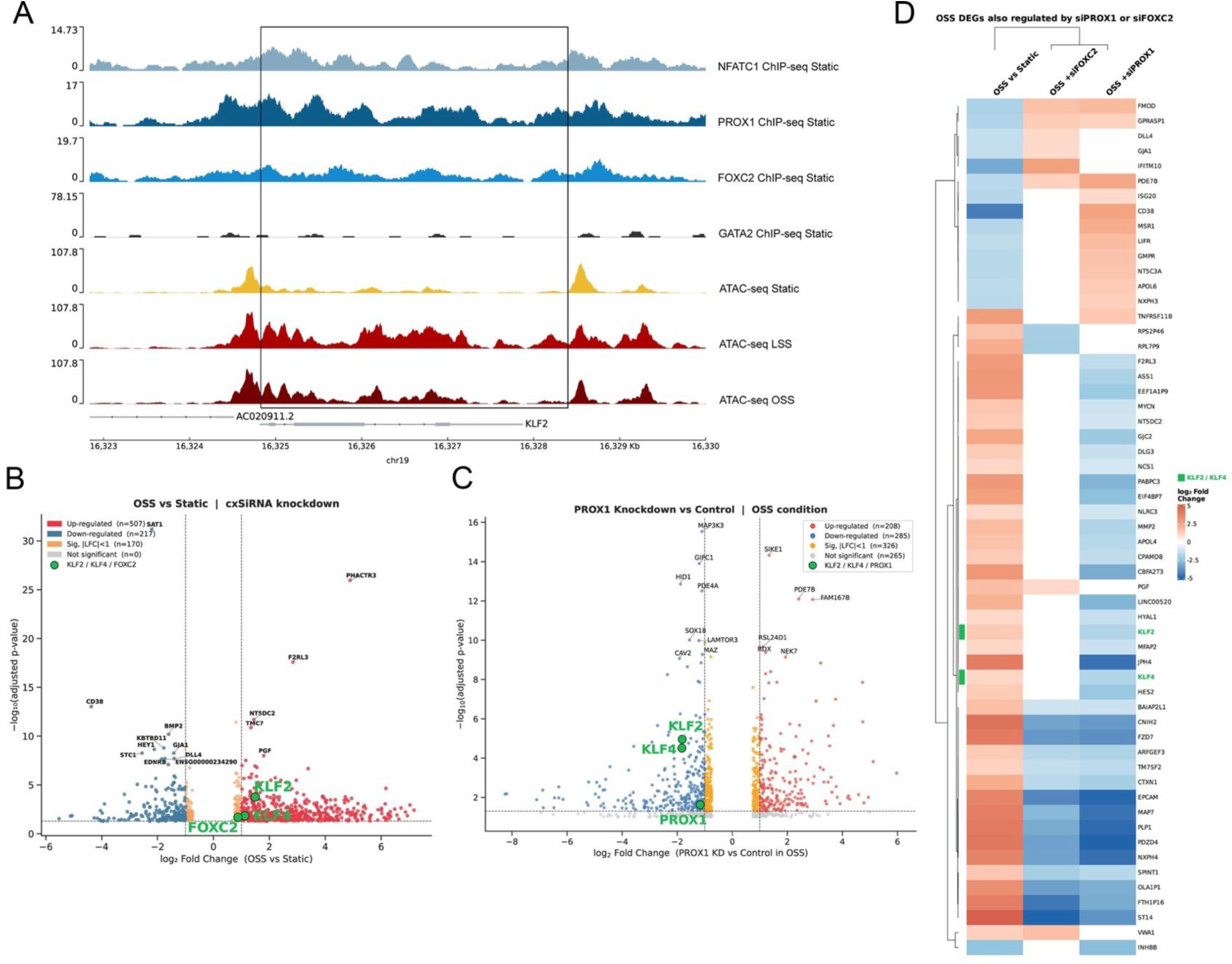
PROX1 and FOXC2 cooperatively and independently regulate shear stress-responsive genes. **(A)** The genomic intervals of the *KLF2* locus. The boxed region shows the chromatin accessibility sites that were increased by both LSS and OSS when compared to the static condition (bottom three rows). The top four rows are from the ChIP-seq dataset and show that PROX1, FOXC2, and NFATC1, but not GATA2, are associated with *KLF2* regulatory elements under static conditions. **(B)** Volcano plot of differentially expressed genes between static and OSS conditions. *KLF2, KLF4*, and *FOXC2* are among the genes upregulated by OSS. **(C)** Volcano plot of differentially expressed genes comparing siControl- and siPROX1-transfected HLECs. *KLF2* and *KLF4* are among the genes that were not upregulated when PROX1 was knocked down. **(D)** Heatmap showing log_2_ fold change (log_2_FC) of 57 genes across three pairwise comparisons: OSS vs static in control siRNA-transfected HLECs; PROX1 knockdown vs control under OSS, and FOXC2 knockdown vs control under OSS. Genes were selected by two criteria: significant differential expression (padj ≤ 0.005, |log_2_FC| ≥ 1.0) in (1) OSS vs static, and (2) at least one knockdown condition. *KLF4* was included regardless of the threshold (OSS vs static values: log_2_FC = 1.12, padj = 0.014), given its established role as a mechanosensitive TF. Color scale: red, upregulated; blue, downregulated; white, no change. **Statistics:** (A) ATAC-seq: n=2 replicates of were performed for each of the three (static, LSS or OSS) conditions; (B-D) Bulk RNA-seq: n=8 control static versus n=5 control OSS; n=2 siPROX1 OSS versus n=5 control OSS; n=2 siFOXC2 OSS versus n=5 control OSS.

To identify upstream regulators of *KLF2*, we intersected ATAC-seq datasets with published ChIP-seq datasets for PROX1, FOXC2, GATA2, and NFATC1. This analysis revealed occupancy of NFATC1, PROX1, and FOXC2 at the *KLF2* locus, whereas no prominent GATA2 binding was detected (**Figure 8A**).

We then tested if PROX1 and FOXC2 regulate *KLF2* expression under OSS, a condition known to induce valve-associated transcriptional programs ^30^. HLECs were transfected with control siRNA, siPROX1, or siFOXC2 and cultured under static or OSS conditions as described previously, followed by bulk RNA-seq to identify downstream responsive genes in an unbiased manner (**Supplementary Tables 12, 13**) ^26,44^.

In control siRNA-transfected HLECs, OSS induced the expression of *KLF2, KLF4,* and *FOXC2* (**Figure 8B**). Differential gene expression analysis further revealed that PROX1 regulates approximately twice as many genes as FOXC2 under these conditions (**Supplementary Figure 5A**). Importantly, OSS failed to upregulate *KLF2* and *KLF4* in siPROX1-transfected HLECs, whereas FOXC2 knockdown had no effect on their expression (**Figure 8C, D** and **Supplementary Figure 5A**). These findings indicate that OSS-induced expression of *KLF2* and *KLF4* is PROX1-dependent but FOXC2-independent.

OSS also induced the expression of genes including *F2RL3, GJC2, JPH4* and *HES2*, and downregulated the expression of *ISG20, CD38,* and *LIFR* in a PROX1-dependent but FOXC2-independent manner (**Figure 8D**). In contrast, OSS-induced upregulation of *EPCAM, FZD7*, and *ARFGEF3*, as well as downregulation of *FMOD* and *PDE7B* required both PROX1 and FOXC2, indicating cooperative regulation. Finally, FOXC2 knockdown selectively impaired the regulation of *DLL4, GJA1,* and *IFITM10*, suggesting that these targets are controlled by FOXC2-dependent but PROX1-independent mechanisms.

To identify the transcriptional programs that are regulated by PROX1 and FOXC2, we compared the differentially expressed genes with ChIP-Atlas 3.0, a publicly available database that catalogs predicted target genes of TFs and chromatin remodelers ^57^. This analysis revealed that siPROX1 preferentially downregulated targets of transcriptional repressors, including EZH2, whereas siFOXC2 affected targets of both transcriptional activators and repressors (**Supplementary Figure 5B**). EZH2 is a histone methyltransferase and the catalytic subunit of Polycomb Repressive Complex 2 (PRC2) ^58^. Several predicted EZH2 targets, including *KLF2, KLF4, SOX18, YAP1, NFATC1* and *NFATC2*, were downregulated by siPROX1 (**Supplementary Figure 5C**). These findings raise the possibility that PROX1 maintains selected OSS-responsive genes by opposing PRC2/EZH2-associated repression.

Together, these findings place PROX1 upstream of FOXC2 in the induction of *KLF2*, *KLF4* and a broad subset of shear-responsive genes, while FOXC2 acts in a gene-selective manner, cooperating with PROX1 at specific targets and independently regulating others.

## DISCUSSION

Lymphatic vascular development is highly sensitive to gene dosage, PROX1 activity and hemodynamic cues, yet how these inputs are integrated has remained unclear. In this study, we identify a dosage-sensitive genetic interaction between GATA2 and FOXC2 that is essential for lymphatic vascular development. While heterozygous loss of either factor alone results in relatively mild defects, combined haploinsufficiency leads to profound abnormalities, including absence of LVs, LVVs, and venous valves, impaired vessel maturation, and reduced survival. These findings establish that GATA2 and FOXC2 function in a partially redundant or buffering manner, particularly during valve morphogenesis and maintenance.

### GATA2 and FOXC2 cooperate to control valve morphogenesis

Our data indicate that valvular endothelial cell specification occurs in *Gata2^+/-^;Foxc2^+/-^* embryos, but they fail to persist and mature into functional valves. This observation places the requirement for GATA2 and FOXC2 cooperation downstream of lineage specification. The severe phenotype observed in compound heterozygotes suggests that these factors provide parallel inputs into a shared developmental program, such that reduction in both lowers transcriptional output below a critical threshold required for valve formation.

### Dosage- and context-dependent regulation of the lymphatic master regulator PROX1

The phenotypic similarity between *Gata2^+/-^;Foxc2^+/-^* and *Prox1^+/-^* mice suggests that PROX1-dependent programs represent a key downstream convergence point. However, our data indicate that this relationship is not linear. While loss-of-function of GATA2 or FOXC2 does not reduce PROX1 expression and may even increase it, GATA2 and FOXC2 overexpression suppresses PROX1 and disrupts lymphatic development ^32^.

These observations are incorporated in **Supplementary Figure 6A**, which proposes that GATA2 and FOXC2 compete with, and thereby constrain, PROX1 activity at a shared regulatory element within the *PROX1* locus. In this framework, PROX1 expression must be maintained within a narrow functional window: insufficient levels impair valve development, whereas excessive expression is also deleterious, consistent with prior experimental evidence ^36,59–61^. GATA2 and FOXC2 may therefore act as rheostatic regulators that prevent inappropriate activation of PROX1-dependent programs.

### Integration with broader mechanotransduction networks

Lymphatic vascular development is regulated by multiple biochemical and mechanosensitive signaling pathways, including Piezo1, VEGF-C, Wnt/β-catenin, YAP/TAZ, BMP9 and S1PR1, all of which have been linked to PROX1, FOXC2 and GATA2 regulation ^14,23,26,32,44,45,62–65^. PROX1 also regulates both GATA2 and FOXC2 in cooperation with Wnt/β-catenin signaling ^45^.

Our findings place PROX1, FOXC2 and GATA2 within this broader framework as integrators of mechanical and biochemical inputs, operating through tightly controlled positive and negative feedback loops to ensure proper lymphatic vascular development. Disruption of this regulatory balance may contribute to lymphatic valve failure and lymphedema.

### PROX1 integrates shear-stress-responsive transcriptional programs

KLF2 and KLF4 are among the best-characterized shear stress-responsive TFs in endothelial cells. In blood vessels, they promote vascular quiescence, maintain vascular tone, and suppress inflammation and thrombosis ^66^. Although their roles in the lymphatic vasculature are less well defined, deletion of *Klf2* alone or in combination with *Klf4* results in severe lymphatic vascular defects ^35,67^. We have determined that PROX1 is required for the induction of *KLF2* and *KLF4*, as well as a broader subset of OSS-responsive genes.

Previous studies have shown that PROX1 can associate with KLF2 to regulate gene expression ^35^. Here, we uncover an additional layer of mechanistic interplay between these factors, demonstrating that PROX1 is required for OSS-mediated upregulation of *KLF2* and *KLF4* in HLECs. These findings support **Supplementary Figure 6B**, in which PROX1 and KLF2 participate in a positive regulatory module. Thus, PROX1 functions not only as a lymphatic identity factor, but also as a central integrator of shear stress-responsive transcriptional programs in LECs.

### FOXC2 modulates gene-specific responses to shear stress

In contrast to the central role of PROX1, FOXC2 regulates distinct subsets of shear-responsive genes. Our data show that FOXC2 is required for the regulation of certain genes independently of PROX1, while others require both factors **Supplementary Figure 6C, D**. FOXC2 regulates a subset of genes independently of PROX1, potentially through interactions with other TFs such as ETS proteins. This is consistent with prior evidence that FOXC2 forms complexes with ETS factors and regulates genes such as *DLL4* and *GJA1* ^52,68,69^.

Together, these models suggest that FOXC2 functions as a context-dependent modulator, shaping the specificity of shear stress responses rather than acting as a global regulator of all shear stress-responsive genes.

### Clinical implications

These findings have implications for understanding Emberger syndrome and other incompletely penetrant vascular disorders. Our results support a dosage-sensitive modifier model for incompletely penetrant lymphatic disease. In patients with GATA2 haploinsufficiency, hematologic malignancies often arise in association with “second hits,” such as monosomy 7, trisomy 8, or acquired mutations in ASXL1 or STAG2 ^70^. Although our data do not demonstrate a second-hit mechanism in patients with Emberger syndrome, they suggest that partial loss of GATA2 can sensitize the lymphatic vasculature to additional genetic or regulatory perturbations. In this context, reduced FOXC2 activity, due to genetic, epigenetic, environmental or hemodynamic factors, represents one such modifier capable of unmasking severe lymphatic vascular disease. In addition, our data raise the possibility that stabilizing FOXC2- or PROX1-dependent transcriptional programs, rather than FOXC2 or GATA2 itself, could mitigate lymphatic dysfunction in susceptible individuals.

### Limitations and future directions

Several limitations must be considered. First, our chromatin analyses relay on integration of our ATAC-seq with previously published ChIP-seq datasets obtained under static conditions. Direct profiling of GATA2, FOXC2, and PROX1 binding under defined shear stress conditions would strengthen mechanistic conclusions. Second, while our GATA2 gain-of-function model supports the model in **Supplementary Figure 6A**, the extent to which PROX1 suppression reflects direct transcriptional repression versus secondary effects on cell survival or differentiation remains to be determined. Third, the proposed models are based on integration of genetic, transcriptional, and chromatin data, and will require targeted perturbations of individual regulatory elements for validation. Finally, this GATA2-PROX1-FOXC2 regulatory network is likely to function in a context-dependent manner, varying across LEC subtypes (capillaries, collecting vessels, and valves) and developmental stages. Elucidating how this network is dynamically configured under physiological and pathological conditions will require further investigation.

In summary, our study defines a dosage-sensitive GATA2-FOXC2 regulatory axis that is essential for lymphatic vascular development. We propose a unified model in which: (1) GATA2 and FOXC2 constrain PROX1 activity, (2) PROX1 integrates shear stress signaling through KLF-dependent programs, and (3) FOXC2 modulates gene-specific transcriptional response in both PROX1-dependent and independent manners. Together, these findings identify PROX1 as a central integrator of shear stress-responsive transcriptional programs and support a dosage-sensitive modifier model in which combined disruption of GATA2- and FOXC2-dependent regulatory inputs causes severe lymphatic vascular disease.

## MATERIALS AND METHODS

### Mice

*Gata2^+/GFP^* (*Gata2^+/-^*) ^71^, *Foxc2^+/-^* ^22^, *Prox1^f/f^* ^72^, Tie2-Cre ^73^, were described previously. The transgenic Gata2^GOF^ model was generated using the strategy described previously ^32,74^. Mice were maintained in C57BL6:NMRI mixed background. All the mice were fed a standard chow diet.

Both male and female mice were used for ex-vivo valve analysis, and similar results were observed for both sexes. The sex of embryos that were used for the study was not determined.

Study Approval: All mice were housed and handled according to the institutional IACUC protocols: OMRF protocols 25-39 and 24-30, and University of Missouri protocol 9797.

### Immunohistochemistry

IHC on cryosections, mesentery and skin were performed according to our previously published protocols ^22,32,45^. A list of primary and secondary antibodies is provided as **Supplementary Table 14**.

### Ex vivo analysis of lymphatic valves

Vessel isolation and cannulation: Mice were anesthetized with ketamine/xylazine (100/10 mg/kg, i.p.) and placed in the prone position on a heated tissue dissection/isolation pad. A popliteal lymphatic vessel containing 3-4 valves was exposed through a superficial incision in the leg, removed and transferred to a Sylgard-coated dissection chamber filled with Krebs-albumin solution for removal of fat and connective tissue. The vessel was then transferred to a 3-ml myograph chamber containing Krebs-albumin solution and cannulated at each end with a glass micropipette (40-50 mm OD tip) and further cleaned while pressurized. The segment was shortened to a single valve for valve function tests, and the remainder of the segment was stored at room temperature for later re-cannulation and study. The chamber, with attached micropipettes, pipette holders and micromanipulators, was transferred to the stage of an inverted microscope. Polyethylene tubing connected the back of each micropipette to low-pressure transducers and a computerized pressure controller ^75^, allowing independent control of inflow (Pin) and outflow (Pout) pressures. Pin and Pout were briefly set to 10 cmH2O at the beginning of each experiment, and the segment was stretched axially to remove longitudinal slack. After returning the pressure to 3 cmH2O, the vessel was allowed to equilibrate in Ca^2+^-free Krebs buffer at 37°C for 20-30 min to eliminate spontaneous contractions that could otherwise interfere with valve function tests. Constant exchange of buffer at a rate of 0.5 mL/min was maintained using a peristaltic pump. Custom LabVIEW programs (National Instruments; Austin, TX) acquired analog data from the pressure transducers at 30 fps simultaneously with vessel inner diameter, which was detected from video images acquired using a Basler firewire camera ^76^. Digital videos of the valve function protocols with embedded pressure data were recorded for additional off-line analyses if needed.

Valve function tests: A 10-mm initial hole was made in the vessel wall near the Pin pipette with a sharply tapered pilot micropipette, which was then removed and replaced with a more gradually tapered servo-null micropipette (tip diameter = 3-5 mm) to measure luminal pressure on the inflow side of the valve (Psn). After insertion, the servo-null micropipette was advanced to seal the hole. The calibration of the servo-null pipette was adjusted as needed after raising Pin and Pout simultaneously between 0.5 and 10 cmH_2_O.

To ensure accurate and consistent measurements of valve back leak, the Pin, Psn, Pout transducers were calibrated before each experiment, and the same pair of cannulation pipettes was used for all experiments to maintain consistent resistances. The maximum value of Psn at Pout=10 cmH_2_O depended on valve resistance and the resistances of the cannulating pipettes and the vessel, but was typically ∼4.7 cmH_2_O.

Pressure back leak through a closed valve was measured by the following procedure. Starting with Pin and Pout = 0.5 cmH_2_O and the valve open, Pout was raised to 10 cmH_2_O, ramp-wise, over a 35-sec period while Pin was held at 0.5 cmH_2_O. After the valve closed, pressure back leak was measured with the servo-null micropipette on the inflow side of the vessel, which could resolve changes as small as ∼0.05 cmH_2_O. The Pout ramp was repeated 3 times. The values of Psn at each Pout level were determined offline using a LabVIEW program by binning the Psn data in 0.5 cmH_2_O Pout intervals.

The adverse pressure gradient (DP, Pout - Pin) required to close an initially open valve. As demonstrated previously ^77,78^, this value increased with increasing vessel diameter. The measurements were therefore made over a wide range of baseline pressures, each of which, in turn, determined the baseline diameter. Starting with the valve open, the output pressure was raised ramp-wise and DP was measured at the instant of valve closure. The test was repeated for baseline pressures 0.1, 0.2, 0.3, 0.5, 1, 2, 3, 5, 8, and 10 cmH_2_O, spanning a range of diameters from ∼40% to 100% of the maximal passive diameter. DP for closure was then plotted against baseline pressure or normalized diameter after the experiment. The highest DP that could be tested was 30 cmH_2_O (equating to a maximum Pout of 40 cmH_2_O when Pin was 10 cmH_2_O) without exceeding the specified safe range of the pressure sensor elements.

Measurements of valve leaflet and gap dimensions: Measurements of leaflet dimensions were made for each valve under brightfield illumination. The side views of the valves used in functional tests did not permit the entire leaflet (the curved insertion paths of the leaflets from their common base to their tips) to be visualized, nor was a measurement of the cusp length usually possible. As an alternative, the best approximations of the leaflet lengths were measured while focusing on the lower surface of the vessel, and then the corresponding approximations of the leaflet lengths on the opposite wall were measured while focusing on the upper surface of the vessel. Valve symmetry was defined as the ratio of the two leaflet lengths.

### Cells and treatments

HLECs were a gift of Dr. Donwong Choi and Young-Kwon Hong (University of Southern California) ^26,34,35^. All experiments were conducted using cells until passage (P) 8. Detailed protocols for culturing HLECs under LSS and OSS are in our previous publications ^44,48^.

### ATAC-seq and computational analysis

ATAC-seq: HLECs were grown under static, OSS or LSS conditions as described above. Cells were detached by trypsinization and pelleted by centrifugation at 500 X G at 4 °C. The supernatant was removed and the cell pellet was resuspended in 500 μl of ice-cold cryopreservation solution containing 50% FBS/40% growth media/10% DMSO. The solutions were transferred to cryotubes on ice, then placed in freezing container and gradually frozen at -80 °C. The frozen tubes were shipped on dry ice to Active Motif (Carlsbad, CA, USA), which performed the downstream steps according to their standard protocols. The experiment was duplicated for each condition.

ATAC-seq data analysis: Sequenced reads were processed with Trim-Galore tool v0.4.4 (https://www.bioinformatics.babraham.ac.uk/projects/trim_galore/) ^79^, potential adapters were removed, and the 3’ ends of reads were quality trimmed with cutadapt (DOI:10.14806/ej.17.1.200) using the quality cutoff of Q20 ^80^. The first 15 bp of each read were clipped to avoid GC bias. Reads were then mapped to the human reference genome (hg38/GRCh38.p13) using Nvidia Parabricks’ FQ2BAM (available on-line: https://docs.nvidia.com/clara/parabricks/v3.5/text/fastq_and_bam_processing.html), that utilize BWA mem v0.7.17-r1188 ^81^ for read mapping and GATK MarkDuplicates ^82^ for duplicated reads identification. Properly paired and uniquely mapped reads were extracted in BAM format with samtools v1.2 (parameter “-q 1 -F 1804,” v1.2) ^83^. After removing mitochondrial DNA reads, the remaining reads were classified into four groups based on fragment size: nucleosome-free and nucleosome reads. The bigwig files were generated using the center 80-bp fragments and scaled to 20 million nucleosome-free reads. Peak calling on the nucleosome-free reads was conducted by MACS2 (v2.1.1.20160309, default parameters with “--extsize 200 -nomodel”) ^84^. To ensure replicability, we first finalized the reproducible peaks for each group as only those that were called with a stringent cutoff (macs2 -q 0.05) in one merged sample and at least called with a lower cutoff (macs2 -q 0.5) in the other merged sample. The reproducible peaks were merged across groups to create a final set of reference chromatin-accessible regions. Peaks were annotated with genes if they overlapped the putative promoter regions, defined as the transcription start site (TSS) and the flanking regions up to 2kbp, with the TSS coordinates being sourced from the Gencode (v31, ^85^) reference annotation. We then counted the nucleosome-free reads from each sample overlapping the reference regions using bedtools (v2.24.0). To elucidate the differentially accessible regions (DARs), we normalized the raw nucleosome-free read counts by trimming the mean using the M-value normalization method. We applied empirical Bayes statistical tests after linear fitting from the voom package (R 3.23, edgeR 3.12.1 and limma 3.26.9). DARs were considered significant when passing absolute log_2_(Fold Change (FC)) > 1 and FDR < 0.05 thresholds. Additionally, as non-differential control regions, we considered genomic intervals that, across all three contrasts, satisfied the following criteria: FC > 1/1.05 and FC < 1.05, p-value > 0.5, and the average log2 CPM (Count Per Million) for both group 1 and group 2 > 0.

Ternary plot based on DARs: Ternary plots were generated for the union of DARs from each comparison. For each genomic interval in the input BED file (0-based, half-open coordinates), we extracted the mean bigWig signal over the full interval using the pyBigWig Python library (stats, type=“mean”, nBins=1; version 0.3.25, available on-line at https://github.com/deeptools/pyBigWig), separately for each of three tracks (one per condition). Intervals with no coverage in a track (missing statistics, None, or non-finite values) were assigned a mean of zero for that track. Peaks for which all three means were zero were excluded from downstream analysis. Compositional coordinates were computed as fi = (mi + p) / Σj(mj + p), where mi is the raw mean for condition i and p ≥ 0 is an optional pseudocount (default of zero was used). Each retained peak was plotted as a single point on a ternary diagram (mpltern; version 1.0.4, available on-line at https://github.com/yuzie007/mpltern; https://doi.org/10.5281/zenodo.3528354), where condition order maps to the top, left, and right vertices (T, L, R). Point color encodes a bivariate Gaussian kernel density estimate (scipy gaussian_kde; version 1.17.1) ^86^: barycentric coordinates were mapped to Cartesian coordinates on the equilateral simplex, density was evaluated at each sample location, and the resulting values were mapped to color. The ternary simplex was partitioned into three regions corresponding to a Voronoi tessellation induced by the three vertices. Region assignment was performed by selecting, for each peak, the condition with the largest compositional component (argmax over fi), with ties resolved by the first maximal index in the order T, L, R (consistent with numpy.argmax). This rule is equivalent to assigning each point to the Voronoi cell of its nearest vertex under the standard equilateral simplex embedding.

Motif Enrichment Analysis (MEA): MEA was performed using HOMER (v5.1) ^87^ via the findMotifsGenome.pl utility. Genomic regions of interest were provided as BED files and analyzed against the specified reference genome build (genome version provided as a parameter). Motif discovery was conducted using the -size given option to preserve the exact input region lengths, with enrichment statistics reported in bits (-bits). Analyses were parallelized across available CPU cores (-p) and executed in local mode (-local 2). The non-differential control regions identified in the DAR analysis were used as the control/background regions. They were supplied to the MEA analysis via the -bg option. Known and custom motif sets were incorporated using the -mknown and -mcheck options, respectively. Output, including identified enriched motifs and associated statistics (e.g., knownResults.txt), was generated per comparison.

### ChIP-seq data processing

Raw reads were downloaded from Gene Expression Omnibus database for the GSE129634 accession number ^25^. Reads were trimmed with Trim Galore (v0.4.4; cutadapt with Q20) ^80^, aligned to hg38/GRCh38.p12 using BWA (v0.7.17-r1198) ^81^, and converted to BAM and filtered with samtools (v1.2) ^83^. Duplicate reads were marked using biobambam2 bamsormadup (v2.0.87) ^88^. Uniquely mapped, non-duplicate reads were retained with samtools (MAPQ ≥1). For QC and fragment size estimation, deduplicated reads were analyzed by strand cross-correlation using SPP (v1.11) ^89^. Next, fragments were extracted to BED format and extended with bedtools (v2.17.0) ^90^, followed by calculating genome-wide coverage using bedtools genomecov in bedGraph mode. Samples were scaled to 15 million sequencing depth, and converted to bigWig format using UCSC bedGraphToBigWig (v4) ^91^.

### Bulk RNA-seq analysis

HLECs were transfected with control siRNA, siPROX1 or siFOXC2 and cultured under static, LSS or OSS condition as described above. Total RNA was extracted from the cells. Subsequently, rRNA was depleted and bulk RNA-seq was performed by Azenta US, Inc (NJ, USA).

## Statistical analyses

In biochemical studies, n refers to the number of times the experiment was performed. For histochemical analysis, n refers to the total number of animals included per group. Statistically significant differences were determined using Student’s unpaired *t-test*, Mann-Whitney test, Chi-square test, one-way ANOVA or 2-way ANOVA with Tukey’s post-hoc tests. Prism software was used for statistical analyses. Data are reported as mean ± SD or mean ± SEM with significance set at p < 0.05. The “n” and “p” values for each experiment are provided in the figure legends.

Gene Set Enrichment Analysis (GSEA) was performed using GSEApy (v1.1.1; https://gseapy.readthedocs.io/en/latest/) in preranked mode ^92^. Genes were ranked using π-value rank ^93^, based on log2(fold-change) ‘ -log10(p-value) values, derived from differential genes or regions analyses, with genes ordered by the absolute value of the ranking metric and duplicate gene symbols collapsed by retaining the highest-ranking entry. As reference gene signatures, the MSigDB ^94–96^ Hallmarks gene set collection (v2023.2.Hs), supplemented by the custom gene set collections, were used. Prior to analysis, gene identifiers in gene rank files, and in-house gene set collections, were harmonized using MSigDB-provided chip files to ensure consistent gene symbol namespaces. GSEA was conducted using a weighted enrichment score, with gene set sizes restricted to a minimum of 15 and a maximum of 500 genes and using 1000 permutations for significance calculations.

## AUTHOR CONTRIBUTIONS

XG, MRM, WR, MvdE, SDZ, CL, MJD and RSS designed and performed experiments and analyzed the results. XG, MJD and RSS wrote the manuscript. HC and AC provided critical input and reagents; all authors provided input in editing the manuscript.

## ACKNOWLEDGEMENTS

We thank Dr. Donwong Choi and Young-Kwon Hong (Beth Israel Deaconess Medical Center, Harvard University) for HLECs, Drs. Brian Coon and Lorin Olson (OMRF) for insightful comments, and Drs. Masayuki Yamamoto (Tohoku University, Japan) and Douglas Engel (University of Michigan) for the *Gata2^+/GFP^* line. This work is supported by OMRF’s institutional funds to RSS, ALSAC (American Lebanese Syrian Associated Charities) fund to CL, NIH/NHLBI (R01HL131652 and R01HL163095 to RSS; R01HL133216 to RSS and HC; R01HL122578 to MJD) and NIH/NIGMS COBRE (P30 GM149376 to XG (PI: Linda Thompson).

## DATA AVAILABILITY

The ATAC-seq and bulk RNA-seq datasets are available from GEO under accession numbers GSE318075 and GSE333179 respectively.

## DECLARATION OF AI USE

The volcano and hierarchical clustering plots in Figure 7 and Supplementary Figure 5 were generated using Biomni.phylo.bio after uploading the spreadsheets containing differentially expressed genes. ChatGPT (version 5.2) was used to assist with editing sections of the manuscript for grammar, clarity and concision.

## Supplementary Figures and Figure Legends

**Supplementary Figure 1.**
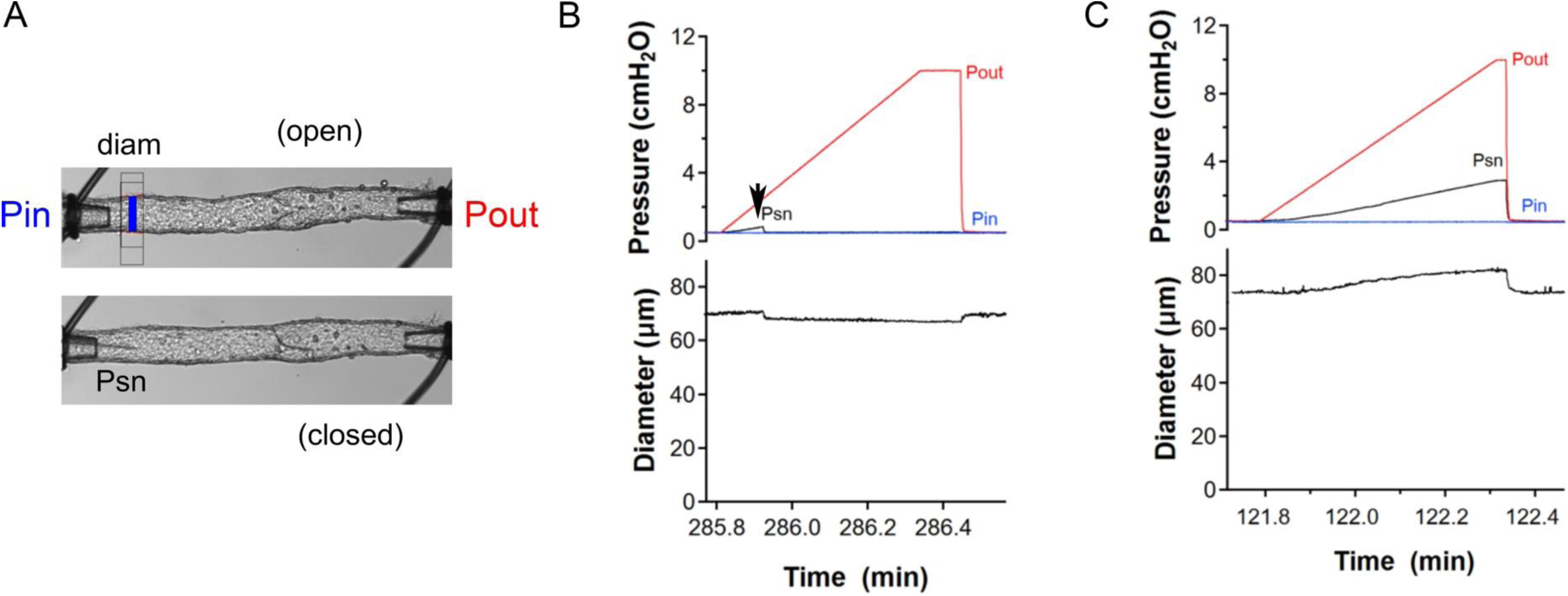
Valve function tests. **A)** Images of a cannulated, pressurized popliteal lymphatic collector containing a single valve, showing the relative position of the valve to the inflow (Pin) and outflow (Pout) pipettes, the diameter tracking site (upper panel), and the placement of the Psn pipette (lower panel) after insertion into the vessel lumen for measurement of intraluminal pressure. Calibration bar = 50 μm. **B**) recordings showing the changes in diameter and the three pressures during a functional test on a normal valve. Pin was held constant at 0.5 cmH_2_O while Pout was elevated, ramp-wise, from 0.5 to 10 cmH_2_O. During the ramp, Psn rose initially when the valve was open, but returned abruptly to 0.5 cmH_2_O when the valve closed (arrow). **C**) Same protocol as in B conducted on an abnormal valve, which closed at arrow but never sealed sufficiently to prevent back leak, such that Psn rose during the Pout ramp from 0.5 to 2.9 cmH_2_O. Diameter at the inflow end of the vessel also increased with increasing Pout.

**Supplementary Figure 2.**
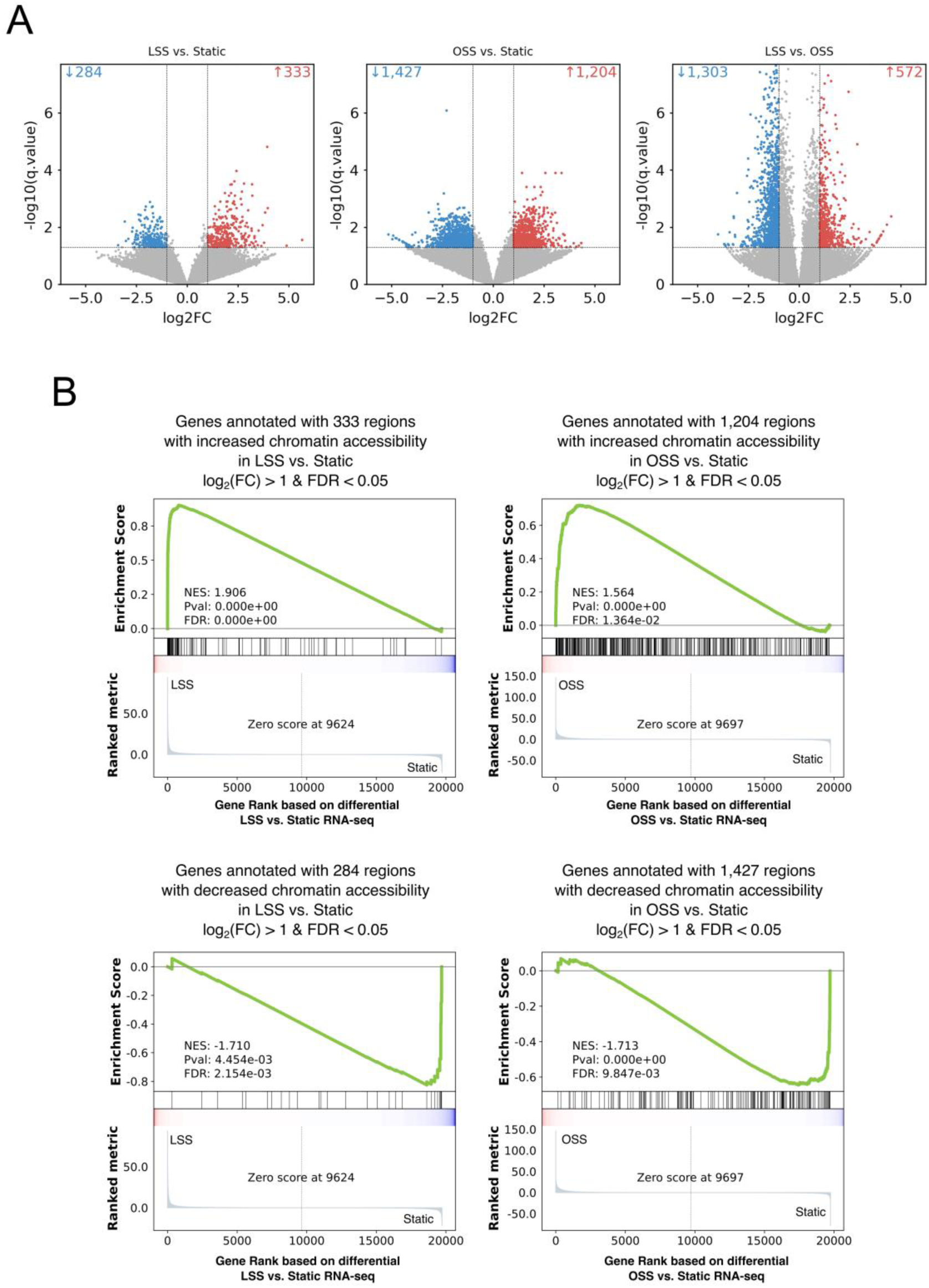
Integration of chromatin accessibility and transcriptional outputs. **(A)** Differential peak calling from ATAC-seq datasets. Volcano plot showing significantly enriched and depleted peaks, with thresholds for statistical significance and fold change indicated. **(B)** Gene set enrichment analysis (GSEA) integrating ATAC-seq and RNA-seq datasets. Gene sets derived from differential chromatin accessibility were compared against ranked RNA-seq expression data to identify concordant regulatory programs. Enriched gene sets highlight coordinated transcriptional and chromatin accessibility regulation. **Statistics:** ATAC-seq: n=2 replicates for each of the three (static, LSS or OSS) conditions; RNA-seq: n=8 static, n=5 OSS and n=3 LSS.

**Supplementary Figure 3:**
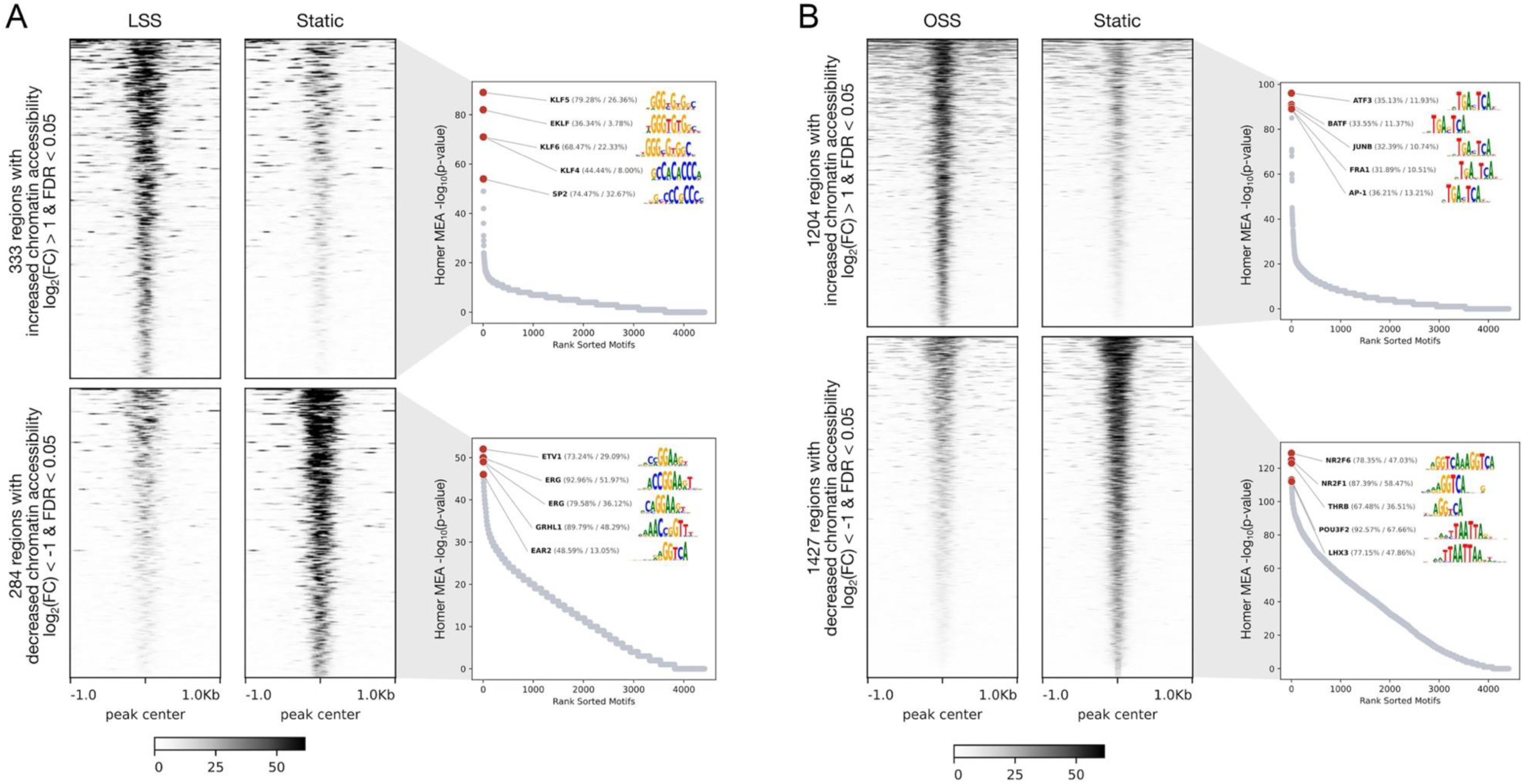
Shear stress alters chromatin accessibility at regions enriched for distinct transcription factor motifs. **(A)** Tornado plots of chromatin accessibility sites that were significantly altered by LSS. TF motif enrichment analysis revealed significant enrichment of KLF motifs and downregulation of ETS motifs in response to LSS. **(B)** Tornado plots of chromatin accessibility sites that were significantly altered by OSS. A significant enrichment of AP-1 motifs and downregulation of nuclear hormone motifs were observed in response to OSS. **Statistics:** n=2 replicates of ATAC-seq were performed for each of the three (static, LSS or OSS) conditions.

**Supplementary Figure 4:**
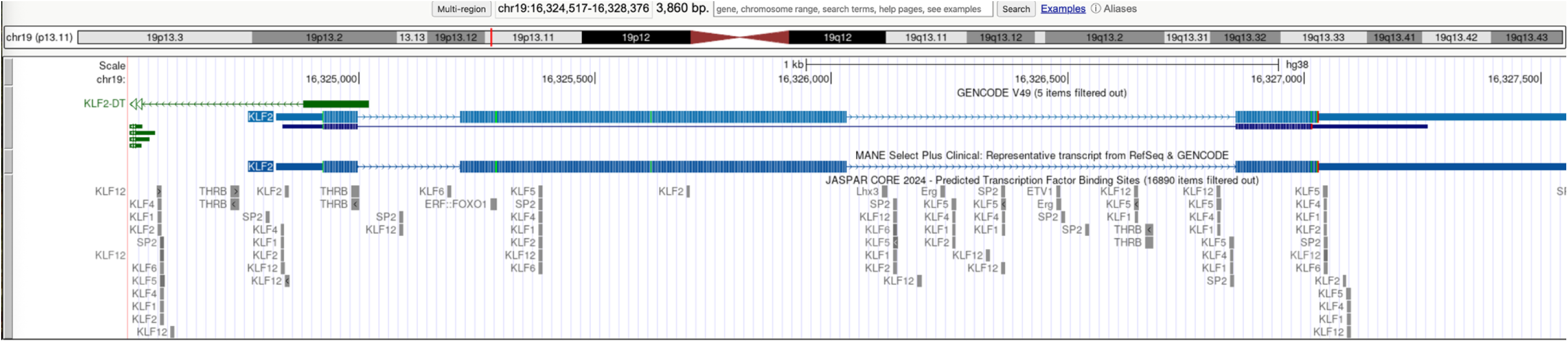
Multiple KLF motifs are observed in the *KLF2* genomic locus. Analysis of the human KLF2 genomic locus using UCSC Genome Browser revealed multiple motifs corresponding to shear stress-sensitive TF families, including KLF, SP2, ETS, and nuclear hormone receptor families. KLF motifs were particularly enriched at this locus.

**Supplementary Figure 5:**
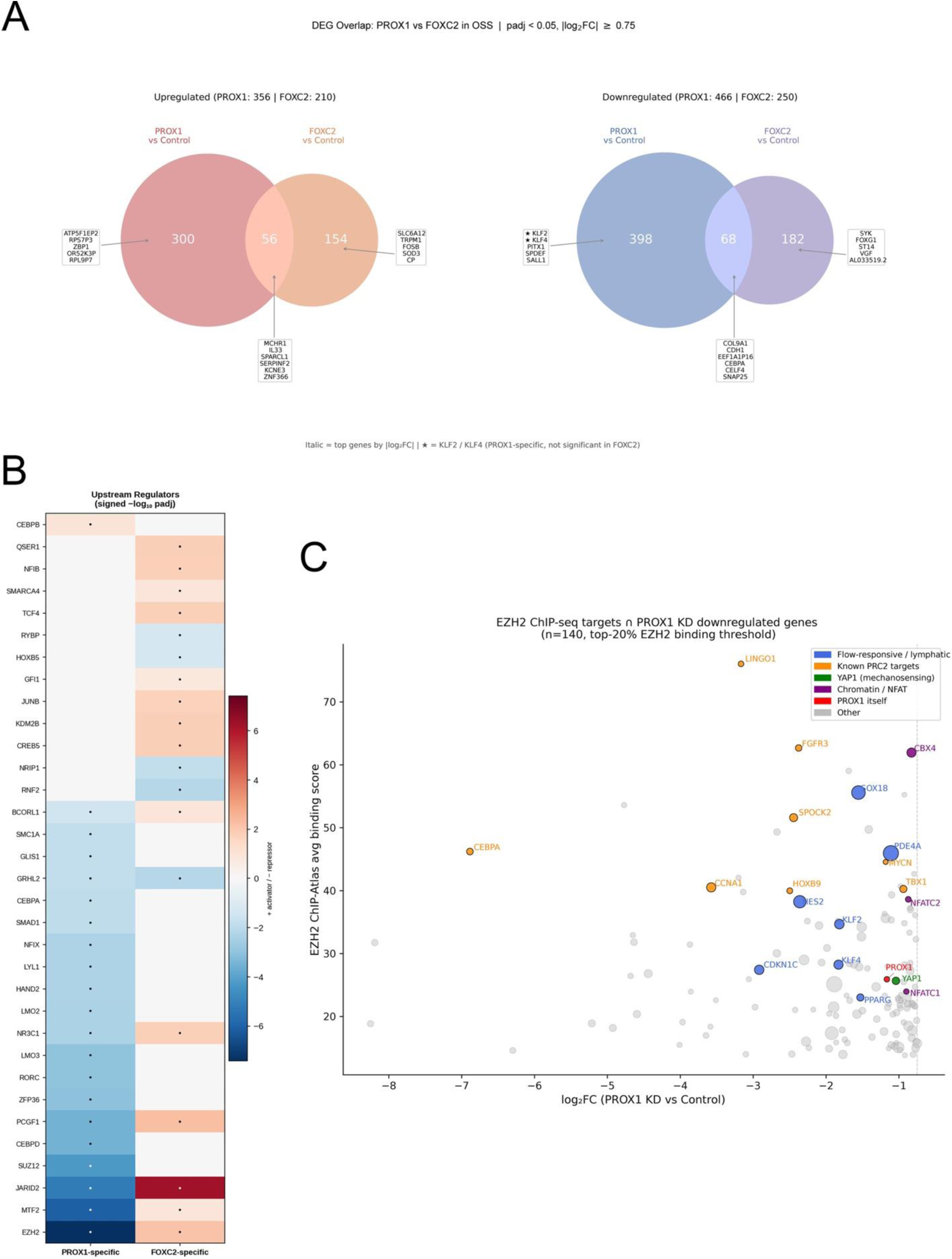
PROX1 and FOXC2 regulate distinct and overlapping OSS-responsive gene programs in HLECs. **(A)** Venn diagrams showing overlap between differentially expressed genes after PROX1 or FOXC2 knockdown in OSS-exposed HLECs. PROX1 knockdown altered more genes than FOXC2 knockdown, with partial overlap between the two groups. Selected PROX1-specific, FOXC2-specific, and shared genes are indicated. Asterisks denote KLF2 and KLF4, which were selectively downregulated by PROX1 knockdown. **(B)** ChIP-Atlas 3.0-based heatmap showing predicted upstream regulators enriched among PROX1-specific and FOXC2-specific differentially expressed genes. PROX1 knockdown preferentially affected candidate targets of transcriptional repressors, including PRC2-associated factors such as EZH2, SUZ12, JARID2, and MTF2, whereas FOXC2 knockdown affected targets of both activators and repressors. **(C)** Scatter plot suggesting EZH2 ChIP-seq targets downregulated following PROX1 knockdown. Highlighted genes include flow-responsive and lymphatic regulatory factors, such as *PROX1, KLF2, KLF4, SOX18*, *YAP1, NFATC1*, and *NFATC2*, suggesting that PROX1 maintains selected OSS-responsive genes by counteracting EZH2/PRC2-associated repression. **Statistics:** n=2 siPROX1 OSS versus n=5 control OSS; n=2 siFOXC2 OSS versus n=5 control OSS.

**Supplementary Figure 6:**
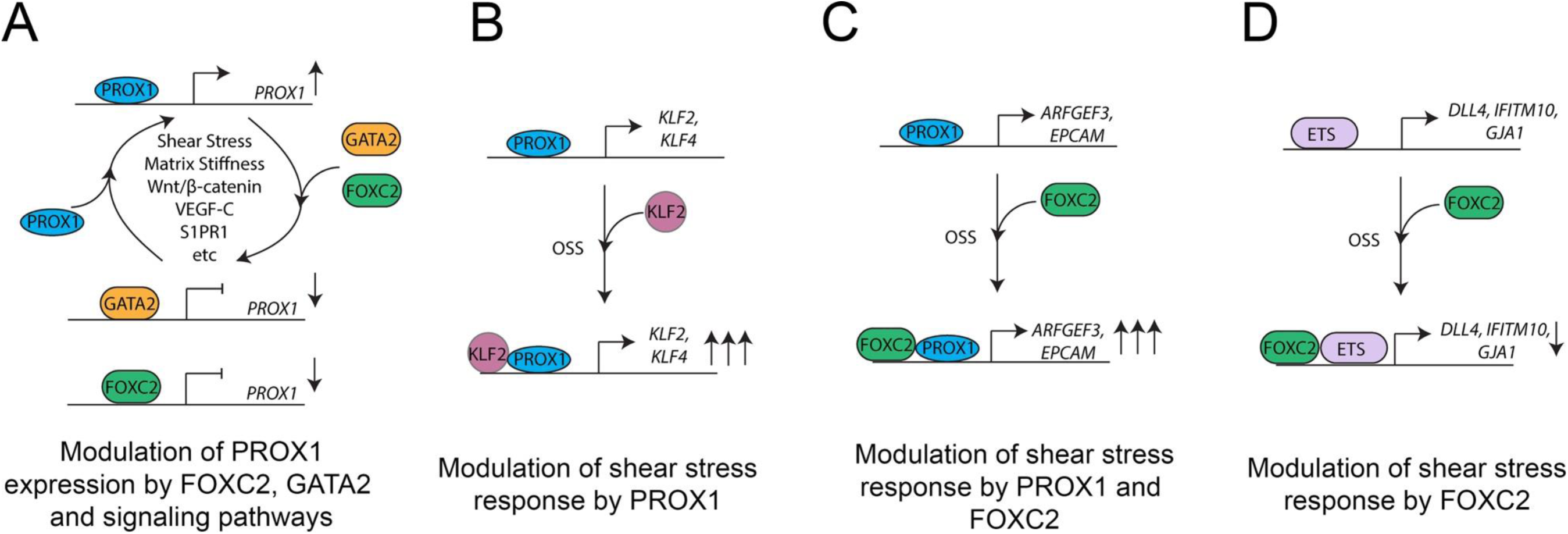
Proposed model for the roles of GATA2, FOXC2 and PROX1 in shear stress response and lymphatic vascular development. (A) GATA2 and FOXC2 compete with PROX1 to associate with the shared regulatory element in the *PROX1* locus. When overexpressed, GATA2 and FOXC2 constrain PROX1 expression. (B) PROX1 and KLF2 form a complex to enhance the expression of *KLF2* and *KLF4*. (C) FOXC2 functions collaboratively with PROX1 to regulate the expression of certain OSS-responsive genes, and (D) FOXC2 functions independently of PROX1 to regulate the expression of certain OSS-responsive genes.

